# ZBTB11 Depletion Targets Metabolic Vulnerabilities in K-Ras Inhibitor Resistant PDAC

**DOI:** 10.1101/2024.05.19.594824

**Authors:** Nathan L. Tran, Jiewei Jiang, Min Ma, Gillian E. Gadbois, Kevin C. M. Gulay, Alyssa Verano, Haowen Zhou, Chun-Teng Huang, David A. Scott, Anne G. Bang, Herve Tiriac, Andrew M. Lowy, Eric S. Wang, Fleur M. Ferguson

**Author notes:** These authors contributed equally (Nathan L. Tran, Jiewei Jiang).

## Abstract

Over 95% of pancreatic ductal adenocarcinomas (PDAC) harbor oncogenic mutations in K-Ras. Upon treatment with K-Ras inhibitors, PDAC cancer cells undergo metabolic reprogramming towards an oxidative phosphorylation-dependent, drug-resistant state. However, direct inhibition of complex I is poorly tolerated in patients due to on-target induction of peripheral neuropathy. In this work, we develop molecular glue degraders against ZBTB11, a C_2_H_2_ zinc finger transcription factor that regulates the nuclear transcription of components of the mitoribosome and electron transport chain. Our ZBTB11 degraders leverage the differences in demand for biogenesis of mitochondrial components between human neurons and rapidly-dividing pancreatic cancer cells, to selectively target the K-Ras inhibitor resistant state in PDAC. Combination treatment of both K-Ras inhibitor-resistant cell lines and multidrug resistant patient-derived organoids resulted in superior anti-cancer activity compared to single agent treatment, while sparing hiPSC-derived neurons. Proteomic and stable isotope tracing studies revealed mitoribosome depletion and impairment of the TCA cycle as key events that mediate this response. Together, this work validates ZBTB11 as a vulnerability in K-Ras inhibitor-resistant PDAC and provides a suite of molecular glue degrader tool compounds to investigate its function.

## INTRODUCTION

Highly aggressive pancreatic ductal adenocarcinomas (PDAC) make up >90% of all pancreatic cancers and are the third leading cause of cancer deaths in the US.^1^ Innovations in surgical management and adjuvant therapy have led to improved outcomes for patients with resectable (Stage 0-IIB) PDAC, who have a 5-year overall survival rate of 21%.^2^ Unfortunately, as early stage PDAC has few symptoms, the majority (>85%) of PDAC patients are diagnosed with unresectable Stage III and Stage IV disease, where the 5-year overall survival remains *below 1%* due to an absence of efficacious anticancer drugs.^2^ Over 95% of PDAC patients harbor tumors driven by activating mutations in K-Ras,^3^ and inhibitors that target oncogenic K-Ras were anticipated to herald paradigm-shifting improvements in patient outcomes. Unfortunately, the initial results in patients treated with drugs targeting K-Ras^G12C^ suggest these benefits may not be realized, due to the rapid onset of K-Ras inhibitor drug resistance.^4^

All of the oncogenic K-Ras mutants found in PDAC are now targetable by at least one clinical-stage inhibitor, and development of both reversible and covalent mutant K-Ras inhibitors is now a highly productive field of research.^5–9^ Currently approved K-Ras inhibitors include sotorasib and adagrasib, which were FDA approved for treatment of K-Ras^G12C^ mutant NSCLC in 2021 and 2022 respectively.^10,11^ Although encouraging responses to sotorasib in PDAC have been reported, resistance swiftly arises.^4^ In a Ph I/II study of patients with K-Ras^G12C^ pancreatic cancer who received at least one previous systemic therapy, an encouraging 21% objective overall response rate and an 84% disease control rate were reported, but this led to a meager 4 month progression-free survival benefit and 5.7 month median duration of response.^4,12^ In both clinical and laboratory studies of K-Ras inhibitor resistance, acquired secondary mutations in K-Ras itself are relatively rare, occurring ∼ 3 – 4% of the time.^13,14^ Resistance to K-Ras inhibition is more commonly mediated by the accumulation of mutations and activation events in other pathways, posing a challenge for the development of second-line K-Ras inhibitors, vertical pathway targeting, or combination therapies once resistance has developed.^15,16^ These data highlight the urgent need to identify and validate new targets that can combat acquired resistance to K-Ras inhibition in PDAC.

Adaptive K-Ras inhibitor resistance is accompanied by significant metabolic reprogramming of cancer cells, including hyperactivation of the PI3K/AKT pathway^17,18^, elevated TCA cycle flux^19^, Myc amplification^20^, and upregulation of mitochondrial biogenesis^21^, culminating in heightened reliance on oxidative phosphorylation (OXPHOS).^21,22^ As K-Ras pathway inhibition impairs glycolysis, forcing cancer cells to rely on OXPHOS for their energetic demands, this mechanism of clinical resistance is expected to impact the entire pharmacological class and is already being reported in laboratory studies of other K-Ras mutant targeting drug candidates.^14,15^ For example, K-Ras^G12D^ is the most common K-Ras mutation in PDAC^3^ and inhibitors that selectively target K-Ras^G12D^ are currently the subject of Ph I clinical trials in PDAC patients (NCT05737706). We recently demonstrated that acquired resistance to the K-Ras^G12D^ inhibitor MRTX1133 in human PDAC cells is associated with feedback activation of ERBB/AKT signaling and enhanced OXPHOS.^14^ Resistance to the mechanistically orthogonal pan K-Ras(ON) inhibitor RM-7977 has been recently reported in murine models of PDAC, and is associated with Myc amplification^20^. While metabolic profiling was not performed in this study, Myc overexpression has been shown to result in increased mitochondrial biogenesis^23^, and over 200 OXPHOS genes are regulated by Myc. These K-Ras inhibitor resistance studies concur with genetic studies in inducible K-Ras^G12D^/p53^+/-^ PDAC murine models, which identified a subpopulation of cancer cells able to overcome genetic ablation of K-Ras^G12D^ to re-seed tumors, and that this subpopulation was highly dependent on OXPHOS.^21^ Pairing oncogenic K-Ras inhibition with OXPHOS inhibition is therefore an attractive therapeutic strategy in PDAC and other Ras addicted malignancies.

Unfortunately, most existing OXPHOS inhibitors have properties limiting their clinical use, including insufficient potency^24^ and poor selectivity^25,26^. Others, such as the mitochondrial complex I inhibitor IACS-010759, display dose-limiting on-target toxicity in the clinic due to their action on actively respiring mitochondria in peripheral neurons.^27^ To identify mechanistically orthogonal approaches to combating OXPHOS-mediated adaptive K-Ras inhibitor resistance, we evaluated public datasets to find targets that met the following four criteria: 1) Upregulated at the protein level during the development of K-Ras inhibitor resistance in PDAC cells at a minimum of one time point.^22^ 2) Essential for cell survival in glycolysis suppression media (low glucose, plus galactose), which forces dependency on OXPHOS.^28^ 3) Not a component of complex I, to identify mechanistically orthogonal targets, with the goal of limiting effects on peripheral neurons and other high-OXPHOS healthy cells.^27^ 4) Chemically tractable via inhibition or targeted protein degradation.^29,30^ This approach identified Zinc Finger and BTB Domain Containing 11 (ZBTB11) as an attractive anti-OXPHOS target for molecular glue degrader development.

ZBTB11 is a zinc finger transcription factor that cooperates with GABPα to regulate the nuclear expression of components of complex I and the mitoribosome, thereby maintaining functional homeostasis in mitochondria.^31–33^ ZBTB11 is transiently upregulated in PDAC cells during the development of K-Ras inhibitor resistance, potentially to sustain increased demands on the biogenesis of mitochondrial components.^22^ Although ZBTB11 is a pan-essential gene^34–36^, we hypothesized that partial loss of ZBTB11 would impair K-Ras inhibitor resistant PDAC fitness but have minimal effects on post-mitotic neurons due to differences in the demands on mitochondrial biogenesis between these cell types.^37^ ZBTB11 contains several C_2_H_2_ zinc finger domains with the CXXCG sequence motif in their beta-hairpin loops, making it potentially chemically targetable via molecular glue degrader-mediated recruitment to the CRL4^CRBN^ E3 ligase complex.^38^ In this study, we validate ZBTB11 as a mechanistically orthogonal target for addressing OXPHOS-dependent drug-resistant states in PDAC, using both genetic and pharmacological approaches.

## RESULTS

### ZBTB11 regulates OXPHOS levels in PDAC cells

To validate that K-Ras inhibitor resistance reliably generates a high-OXPHOS state, and to develop in-house PDAC models, we cultured 3 independent K-Ras^G12C^ MIA PaCa-2 lines with escalating sublethal concentrations of sotorasib (AMG-510) to generate sotorasib-resistant cell lines R1, R2 and R3 (Fig. 1A, Fig. S1A). We characterized the cellular metabolic state using Seahorse bioenergetic analysis, where a mitochondrial stress test revealed increased basal and maximal oxidative phosphorylation in the resistant cells compared to parental cell controls, as well as increased extracellular acidification rate (ECAR) resulting in overall higher concentrations of ATP produced per minute per cell (Fig. 1B-C). We performed Seahorse bioenergetic analysis on our previously reported K-Ras^G12D^ MRTXR SUIT2 lines^14^ which are resistant to MRTX1133 and compared them to parental SUIT2 cells (Fig. S2A-C). Here we also observed increased basal and maximal oxidative phosphorylation, and overall higher concentrations of ATP produced per minute per cell in MRTXR cells. Finally, we performed mass-spectrometry based metabolomics following ^13^C glucose or ^13^C glutamine labeling to identify differences between parental and resistant MIA PaCa-2 cells (Fig. S1B, Dataset 1) and parental and resistant SUIT2 cells (Fig. S2D, Dataset 1). Metabolite set enrichment analysis identified changes in diverse metabolic pathways including branched chain amino acid biosynthesis/degradation, glycolysis/gluconeogenesis, and the TCA cycle. These data highlight the breadth of metabolites and pathways altered during the development of resistance, that culminate in the observed increases in OXPHOS.

**Figure 1.**
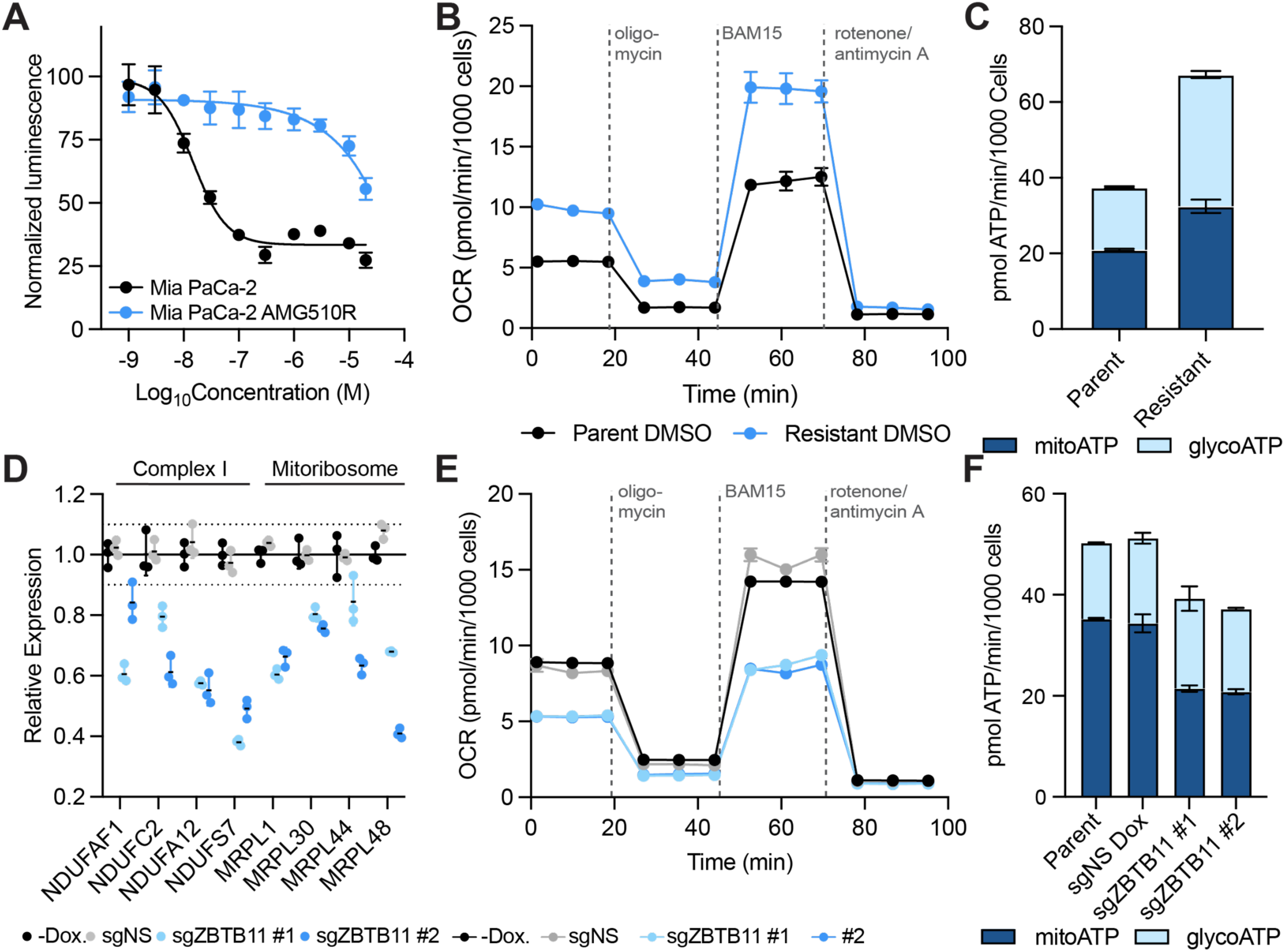
| OXPHOS is upregulated in K-Ras inhibitor resistant PDAC cells, and countered by ZBTB11 knockdown. A. Sotorasib/AMG-510-resistant MIA PaCa-2 cells are 100-fold less sensitive to sotorasib/AMG-510 than parental MIA PaCa-2 cells. Cells were treated with the indicated dose of sotorasib for 72 hrs, and viability was evaluated by CellTiterGlo®. Data is depicted as the average +/− standard deviation (S.D.) of *n* = 3 biological replicates and is normalized to DMSO vehicle treatment controls. See also Fig. S1A. B. Sotorasib-resistant MIA PaCa-2 cells perform higher basal and maximal levels of OXPHOS than parental MIA PaCa-2 cells. C. Sotorasib-resistant MIA PaCa-2 cells are more energetic than parental MIA PaCa-2 cells. D. Expression of genes regulated by ZBTB11 decreases following CRISPRi-mediated knockdown. Cells were treated with 500 ng/ml of doxycycline for 72 hrs to induce Cas9 expression, and mRNA levels were quantified by RT-qPCR. Data is depicted as the average +/− S.D. of *n* = 3 biological replicates and is normalized to H_2_O vehicle treatment controls. See also Fig. S2. E. ZBTB11 knockdown reduces rates of OXPHOS. Cells were treated with 500 ng/ml of doxycycline for 72 hrs to induce Cas9 expression. F. ZBTB11 knockdown reduces ATP production by OXPHOS. B-C, E-F. Cellular respiration rates were evaluated in a mitochondrial stress test using a Seahorse analyzer. Oxygen Consumption Rates (OCR) and Extracellular Acidification Rates (ECAR) from the mitochondrial stress test were used to calculate ATP production rates. Data is depicted as the average +/− S.D. of *n* = 2 biological replicates with *n* = 3 technical replicates each.

ZBTB11 regulates nuclear transcription of components of the mitoribosome and complex I in murine embryonic stem cells and HEK293 cells^33^ to modulate OXPHOS, but its functions in PDAC have not been characterized. To evaluate if human ZBTB11 regulates these genes, we developed inducible ZBTB11 CRISPRi cell lines to evaluate effects of ZBTB11 depletion genetically, confirming knockdown by immunoblot and qPCR (Fig. S3A-B, D). We selected 4 representative mitoribosome genes (MRPL48, MRPL44, MRPL1, MRPL30) and 4 complex I genes (NDUF37, NDUFA12, NDUFC2, NDUFAF1) reported to be regulated by murine ZBTB11^33^, and performed qPCR to quantify their mRNA levels 1 and 3 days after hZBTB11 CRISPRi knockdown. We observed reduction of the mitoribosome mRNA set 24 hrs after dCas9-KRAB induction, and reduction of the complex I mRNA set after 3 days (Fig. 1D, Fig. S3C, E). To test if ZBTB11 depletion leads to reduced OXPHOS, we performed Seahorse bioenergetic analysis and observed the expected reduction in oxygen consumption rate (OCR), with no change to the ECAR 24 hrs following dCas9-KRAB induction (Fig. 1E-F). We conclude that ZBTB11 is a transcriptional regulator of mitochondrial genes and OXPHOS in K-Ras mutant PDAC, consistent with its reported role in other organisms and cell types.

### Development of a CRBN-recruiting molecular glue degrader of ZBTB11

Although ZBTB11 harbors the CXXCG beta-hairpin motif found in many CRBN molecular glue targets, no CRBN-recruiting degraders of ZBTB11 have been reported.^38^ To enable high-throughput quantification of ZBTB11 protein levels in 384 well plates, we used CRISPR/cas9 to knock in a HiBiT tag at the c-terminus of the endogenous ZBTB11 locus and used this reporter system to profile a proprietary library of CRBN-binding molecular glue candidates (Fig. 2A). We identified several multi-targeted hits from the same chemical series that were able to deplete ZBTB11 by greater than 30% (Fig. 2B-C). Medicinal chemistry optimization (to be reported elsewhere), culminated in the identification of JWJ-01-306 (Fig. 2C) which degrades ZBTB11 up to a D_max, 10 µM_ of 60% by HiBiT quantification, and up to a D_max, 10 µM_ of 90% by immunoblot (Fig. 2D, Fig. S4B). We confirmed rescue of JWJ-01-306 mediated ZBTB11 degradation using pre-treatment with proteasomal inhibitor carfilzomib, NAE1 inhibitor MLN4924, and by competition with the CRBN binder lenalidomide (Fig. 2E).^39^

**Figure 2.**
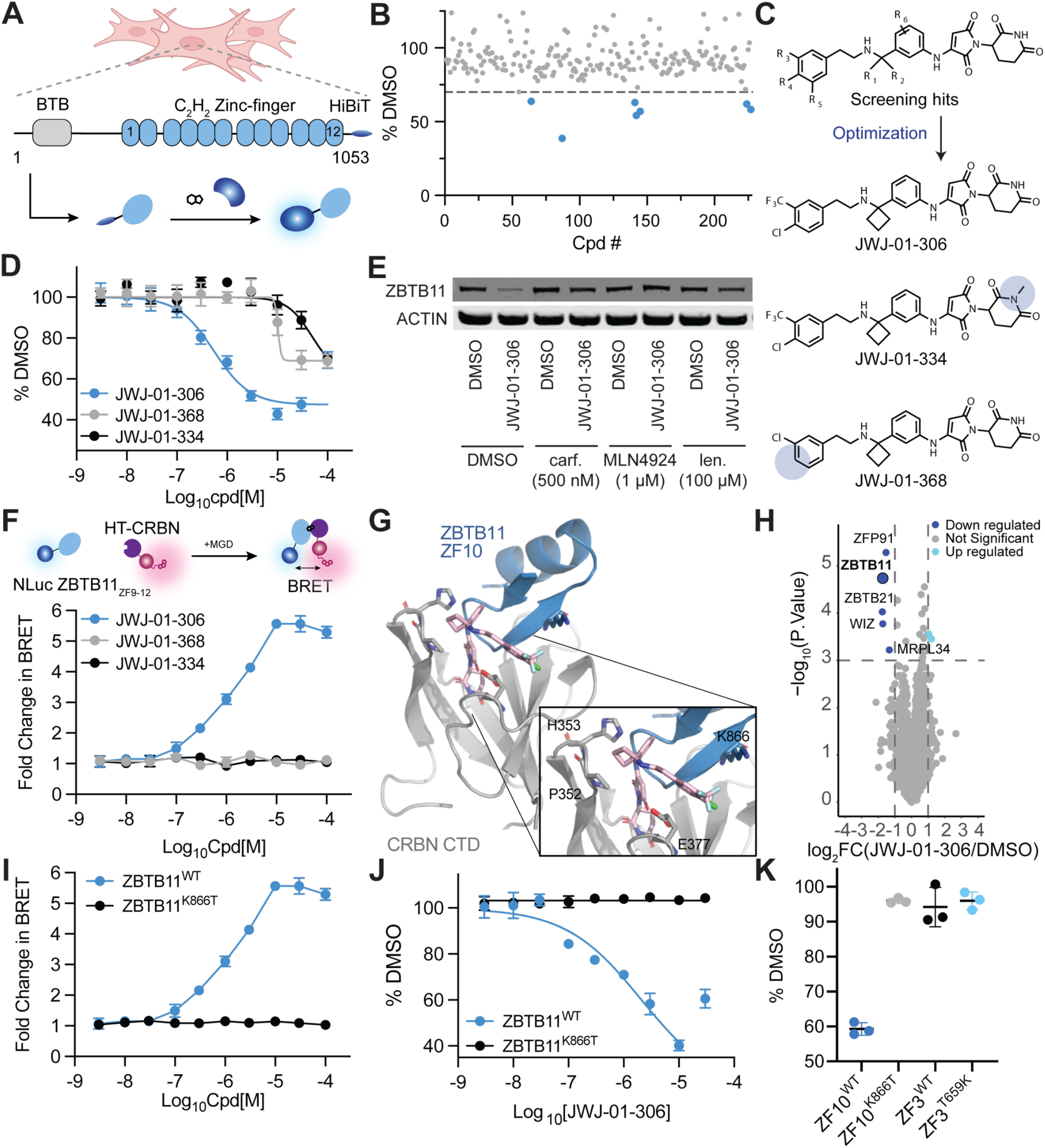
| Development of a ZBTB11 molecular glue degrader. A. Depiction of ZBTB11 domains and the ZBTB11-HiBiT assay. B. Screening of glutarimide-containing analogues in the ZBTB11-HiBiT assay identifies candidate ZBTB11 degraders. MOLT-4 ZBTB11-HiBiT knock-in cells were treated with 10 µM compound for 8 hrs. Data is depicted as the average of *n* = 3 biological replicates and is normalized to DMSO vehicle treatment control. C. Chemical structure of screening hits and optimized ZBTB11 degrader and negative controls. D. JWJ-01-306, but not negative controls, degrades ZBTB11-HiBiT in MIA PaCa-2 knock-in cells. Cells were treated for 5 hrs with the indicated compound. E. Mechanism-based controls rescue ZBTB11-HiBiT degradation in MIA PaCa-2 knock-in cells. Cells were pre-treated with carfilzomib, MLN4924, or lenalidomide for 1 hr prior to treatment with the indicated compound for 5 hrs. Protein levels were quantified by western blot. Depicted blots are representative of *n* = 3 independent experiments. Uncropped blots found in Source Data. F. JWJ-01-306, but not negative controls, induces CRBN:degrader:ZBTB11 complex in NanoBRET™ ternary complex assay. A MOLT-4 cell line with stable expression of the indicated constructs was generated. Cells were pre-treated with carfilzomib and the HaloTag® NanoBRET™ 618 Ligand for 1 hr prior to treatment with the indicated compound for 5 hrs. G. Computationally generated model of the CRBN:JWJ-01-306:ZBTB11 ZF10 complex reveals binding mode and key residues for complex stabilization. Top ten scoring structure clusters found in Source Data. H. Global proteomics analysis of MIA PaCa-2 cells treated with 10 µM JWJ-01-306 for 5 hrs. Samples were prepared as *n* = 3 biological replicates. Full datasets in Dataset 2. I. ZBTB11 K866T ablates ternary complex formation. MOLT-4 cell lines with stable expression of the indicated constructs were generated and used in the NanoBRET™ ternary complex assay. J. ZBTB11 K866T rescues JWJ-01-306-mediated degradation. MIA PaCa-2 cell lines with stable expression of the indicated constructs were generated. Cells were treated with JWJ-01-306 for 5 hrs. K. K866 is necessary but not sufficient for ZBTB11 degradation. MOLT-4 cell lines with stable expression of the indicated constructs were generated. Cells were treated with 1 µM screening hit compound ALV-05-184 for 5 hrs. D, F, I-K. Data is depicted as the average +/− S.D. of *n* = 3 biological replicates and is normalized to DMSO vehicle treatment control.

Molecular glue degraders depend on the formation of ternary complexes for their cellular activity. To measure intracellular ternary complex between our molecules, ZBTB11, and CRBN, we developed a NanoBRET assay, and used it to confirm that JWJ-01-306 promotes a CRBN:molecular glue:ZBTB11 ternary complex (Fig. 2F). To enable structure-based design of improved ZBTB11 degraders and controls, we established ternary complex molecular modeling protocols (Fig. 2G). ZBTB11 contains 12 C_2_H_2_ zinc finger domains, 7 of which harbor a CXXCG motif (Fig. S4F). We generated NLuc tagged ZF constructs and quantified the change in individual ZF-NLuc fusion protein levels upon treatment with ZBTB11 degrader molecules, to elucidate that zinc finger 10 (ZF10) contains the primary CRBN/JWJ degron (Fig. S4C-E). Next, we generated an AlphaFold2^40^ model of ZBTB11 ZF10 and used this structure to perform Rosetta protein-complex docking to JWJ-01-306-bound CRBN.^41^ Manual inspection of the clusters revealed a binding mode consistent with previously reported cryo-EM structures of CRBN-binding molecular glue degraders.^42^ The glutarimide binds in the tri-tryptophan pocket of CRBN where it makes two H-bonds to H378 and one H-bond to the backbone amide of S379 (Fig. 2G). The aniline ring engages in an edge-to-face π interaction with H353 of CRBN, and 3-substitution orients the benzylic cyclobutane ring towards the interface of CRBN and ZBTB11 ZF10, making hydrophobic contact with CRBN P352. Finally, ZBTB11 residue K866 and CRBN residue E377 form a cation-π-anion stacking interaction with the 4-Cl, 3-CF_3_ substituted phenyl ring in JWJ degraders that bridges the ternary complex.^43^ These interactions agree with trends in the ZBTB11 degrader structure activity relationships.

To aid interpretation of phenotypic data and confirm observed effects are on-target, we developed two negative control compounds: JWJ-01-334 and JWJ-01-368. To rescue all CRBN-dependent pharmacology, we synthesized JWJ-01-334, which contains an N-Methyl glutarimide that prevents binding to CRBN^44^, ZBTB11 ternary complex formation, and ZBTB11 degradation (Fig. 2D, F, Fig. S4A). To rescue ZBTB11-dependent pharmacology, we synthesized JWJ-01-368, which lacks the aromatic CF_3_ substituent crucial for achieving the cation-π-anion interaction that promotes CRBN:JWJ:ZBTB11 ternary complex formation. In live cells, JWJ-01-368 efficiently binds CRBN (Fig. S4A) but fails to promote both ZBTB11 ternary complex formation (Fig. 2F) and ZBTB11 degradation (Fig. 2D).

We evaluated the selectivity of JWJ-01-306 and JWJ-01-368 in MIA PaCa-2 cells by global proteomics analysis, following a 5 hr treatment with 10 µM compound (Fig. 2H and Fig. S5A, Dataset 2). In JWJ-01-306 treated cells, we observed potent ZBTB11 downregulation, and off-target degradation of three related C_2_H_2_ zinc finger transcription factors: ZFP91, ZBTB21 and WIZ. In JWJ-01-368 treated cells, we did not observe ZBTB11 degradation, indicating JWJ-01-368 is a suitable control compound for rescue of ZBTB11-specific pharmacology (Fig. S5A, Dataset 2). Next, as common CRBN-recruiting molecular glue degrader off-target IKZF1 is not expressed in PDAC, we performed follow-up screening in a Jurkat IKZF1-HiBiT cell line. Here, JWJ-01-306 and JWJ-01-368 showed potent activity, indicating they would not be selective in certain lymphoid or myeloid cell lines and should be used with caution in that context (Fig. S4G). Finally, to confirm selectivity against the anti-tumor target GSPT1^45^, we performed follow-up screening in a Jurkat GSPT1-HiBiT cell line, where all 3 compounds were inactive (Fig. S4H).

To account for pharmacology that may be mediated by off-target activities of JWJ-01-306, we sought to design a degradation-resistant mutant for use as a genetic control in rescue experiments. From our structural models, we predicted that K866 made critical contributions to ternary complex formation between CRBN and ZBTB11 ZF10 (Fig. 2G). In contrast, ZBTB11 ZF3, which has a CXXCG motif but a threonine (T659) in the equivalent position to K866, was not degraded as an NLuc fusion by our molecules (Fig. 2K). We performed site-directed mutagenesis to generate ZBTB11^K866T^, which ablated both ternary complex formation and degradation of full length ZBTB11 by JWJ-01-306 (Fig. 2I-J). To examine if a lysine in the K866 position alone can promote degradation by JWJ-01-306, we mutated ZF3 T659 to generate ZF3^T659K^. This construct remained recalcitrant to degradation, indicating that K866 is necessary but not sufficient for degradation by our compounds and highlighting the importance of the additional contacts made between CRBN and ZBTB11 ZF10 in the ternary complex that drive the specificity of our degrader proteome-wide (Fig. 2H, K).

To evaluate the kinetics of JWJ-01-306 mediated degradation, we performed time-course studies which indicated that the majority of the observed degradation occurs within 2 hrs (Fig. S5B). Next, we evaluated the rebound rate of ZBTB11 levels following compound washout. Here, we observed no rebound in ZBTB11 levels over a 24 hr time period, although we cannot completely rule out insufficient washout as a confounding factor (Fig. S5C). Finally, we performed pharmacokinetic analysis in C57BL/6 mice dosed via intraperitoneal (IP) injection with 10 mg/kg JWJ-01-306.HCl. Here, we observed a half-life of 91 minutes and a C_max_ of 1.67 µM (Fig. S5D, Dataset 3). Plasma protein binding analysis revealed that JWJ-01-306 is over 99% protein bound (Dataset 3). Taken together, these data indicate that the free drug levels of JWJ-01-306 *in vivo* are unlikely to be sufficient to support ZBTB11 degradation at reasonable dosing regimens, and that further chemistry optimization is needed to improve the stability and physicochemical properties of JWJ-01-306. Therefore, JWJ-01-306 is best suited as a cell-based tool compound for target validation and mechanistic studies.

### ZBTB11 degradation impairs proliferation of K-Ras inhibitor resistant PDAC cells

To determine the effectiveness of our ZBTB11 degraders for targeting K-Ras inhibitor-resistant PDAC, we performed cell proliferation assays via cell counting over time (MIA PaCa-2, 14d) and live cell brightfield-imaging (SUIT2, 6d) (Fig. 3A-B). As expected, proliferation of parental MIA PaCa-2 cells was completely impaired by high concentrations of sotorasib. In these highly glycolytic cells,^46^ single agent JWJ-01-306 treatment had only mild effects on proliferation over a 2-week time period, comparable to the negative control JWJ-01-368 (Fig. 3A). K-Ras inhibitor-resistant MIA PaCa-2 cells showed significantly impaired proliferation only in the presence of sotorasib + JWJ-01-306 (Fig. 3A), but not sotorasib + JWJ-01-368. In SUIT2, proliferation of parental cells was completely impaired by high concentrations (500 nM) of MRTX1133 and by JWJ-01-306 as a single agent within 5 days (10 µM, Fig. 3B). Gratifyingly, cell growth in MRTX1133-resistant SUIT2 lines was blocked by JWJ-01-306 single agent treatment and MRTX1133 + JWJ-01-306, but not MRTX1133 alone or MRTX1133 + JWJ-01-368 (Fig. 3B). These data demonstrate that JWJ-01-306 can counter K-Ras inhibitor resistance in multiple types of mutant K-Ras-driven PDAC and may also have single agent antiproliferative activity in some PDAC contexts.

**Figure 3.**
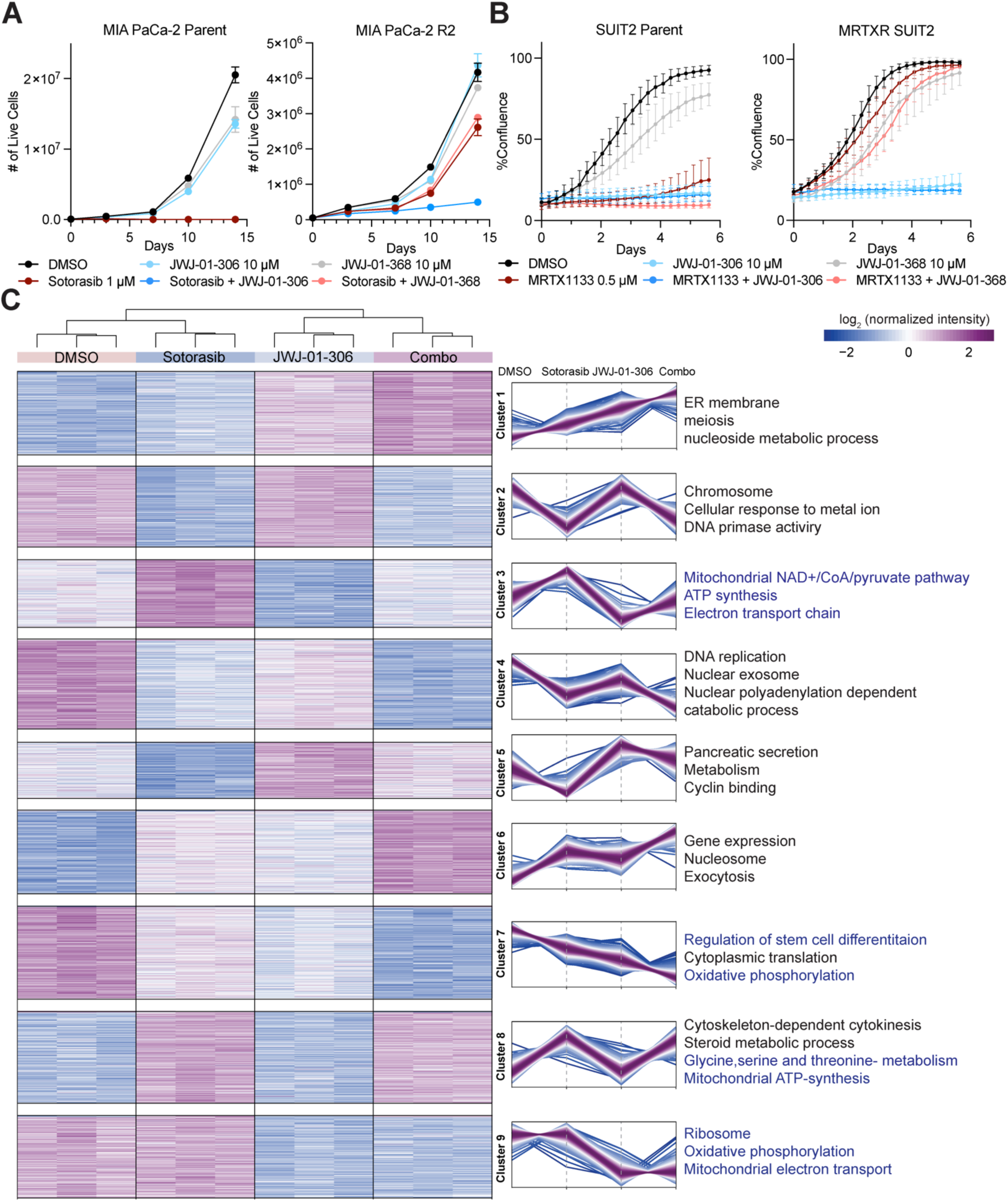
| JWJ-01-306 combination treatment overcomes acquired resistance to K-Ras inhibitors in PDAC via metabolic pathway reprogramming. A. JWJ-01-306 synergizes with sotorasib to inhibit proliferation of sotorasib-resistant MIA PaCa-2 cells. Cells were treated with the indicated compounds and cell numbers were measured using trypan blue exclusion. Data is depicted as the average +/− S.D. of *n* = 3 biological replicates. B. JWJ-01-306 effectively inhibits proliferation of SUIT2 cells. Cells were treated with the indicated compounds and cell confluence was measured using a Cellcyte X live cell analyzer. Data is depicted as the average +/− S.D. of *n* = 3 biological replicates with *n* = 4 technical replicates each. C. Sotorasib-resistant MIA PaCa-2 cells were treated with DMSO, 1 µM sotorasib, 10 µM JWJ-01-306, or 1 µM sotorasib + 10 µM JWJ-01-306 (combo) for 72 hrs followed by global proteomics analysis, *n =* 3 biological replicates per condition. Protein abundance changes were analyzed by one-way ANOVA test (FDR < 0.05) and clustered using K-nearest neighbors clustering. Each line within the clusters is color-coded according to its distance from the cluster center, ranging from purple (close) to light blue (far). Proteins in each cluster were then evaluated for pathway enrichment using GO and KEGG pathway analysis. Representative pathways are shown and pathways associated with ZBTB11 function are highlighted in blue text. Full datasets in Dataset 4.

### ZBTB11 degradation reprograms cellular metabolism to reduce OXPHOS and the TCA cycle

We next sought to mechanistically interrogate the metabolic, transcriptomic and proteomic changes induced by JWJ-01-306 in the context of K-Ras inhibitor-resistant PDAC. To identify how both K-Ras inhibitors and ZBTB11 degraders affect the cellular state at phenotypically relevant time points, we performed global proteomics analysis of sotorasib-resistant MIA PaCa-2 cells treated with DMSO, 1 µM sotorasib (resistant cell media), 10 µM JWJ-01-306 and 1 µM sotorasib + 10 µM JWJ-01-306 for 3 and 5 days (Fig. 3C, Fig. S6, Fig. S7, Dataset 4). We compared the proteomic changes between treatment groups and performed K-nearest neighbor clustering analysis to identify protein groups with the same changes in protein abundance in response to each treatment condition (Fig. 3C, Fig. S6, Dataset 4). Next, we performed gene ontology analysis to identify the biological pathways enriched within each protein cluster. Proteins downregulated by JWJ-01-306 treatment correspond to mitochondrial, oxidative phosphorylation, mitoribosome, and differentiation pathways (Fig. 3C cluster 3, 7 and 9, Fig. S6 cluster 4). Protein sets primarily responsive to sotorasib include upregulation of mitochondrial ATP synthesis, and changes to glycine, serine, and threonine metabolism pathway proteins, consistent with the observed metabolic and metabolomic differences we observed between parental and sotorasib-resistant cell lines (Fig. 3C cluster 8, Fig. S6 cluster 5). Finally, clusters representing proteins that change only in response to the combination treatment are enriched for nucleoside metabolism, ER membrane, and gene expression (upregulated), and DNA replication (downregulated, Fig. 3C cluster 4). To further elucidate the effects of ZBTB11 depletion on mitochondrial proteins, we evaluated proteins downregulated in JWJ-01-306 treatment conditions and compared them to the MitoCarta 3.0 database of mitochondrial proteins (Fig. S7A). We observed significant overlap (592 proteins day 5, 554 proteins day 3), in each condition with mitochondrial proteins. We confirmed ZBTB11 is downregulated in JWJ-01-306 treated samples (Fig. S7B) alongside a subset of ZBTB11 downstream proteins (NDUFS7, NDUFA12, NDUFC2, NDUFAF1, MRPL48, MRPL44, MRPL1, MRPL30, Fig S7B). K-Ras levels were also reduced in the 5 d samples in response to JWJ-01-306 but not sotorasib (Fig. S7B).

To evaluate the impact of ZBTB11 degradation and consequent downstream proteome changes on OXPHOS levels, we treated parental and sotorasib-resistant MIA PaCa-2 cells with 10 µM JWJ-01-306, JWJ-01-334 or JWJ-01-368 and performed a mitochondrial stress test at the 24 hr time point (Fig. 4A, Fig. S8A). We observed significant reductions in basal and maximal oxidative phosphorylation in cells treated with JWJ-01-306 but not JWJ-01-334 or JWJ-01-368. We also observed changes in the metabolic index (mitoATP/glycoATP ratio) in cells treated with JWJ-01-306 but not JWJ-01-334 or JWJ-01-368, reflecting the greater reduction in OXPHOS (Fig. 4B, Fig. S8B). Finally, we repeated these experiments in parental and MRTX1133-resistant SUIT2 cells, and also observed JWJ-01-306-dependent reductions in basal, maximal, and mitoATP/glycoATP ratios, that were rescued by the control compounds (Fig. 4C-D, Fig. S8D-E). To confirm these effects are specific to ZBTB11 degradation, we stably expressed full length ZBTB11^WT^ or ZBTB11^K866T^ in sotorasib-resistant MIA PaCa-2 cells, followed by treatment with 10 µM JWJ-01-306, and performed a mitochondrial stress test after 24 hrs of drug treatment (Fig. 4E). We observed complete rescue of oxidative phosphorylation and restoration of the metabolic index by the ZBTB11^K866T^ mutant relative to WT control (Fig. 4E-F).

**Figure 4.**
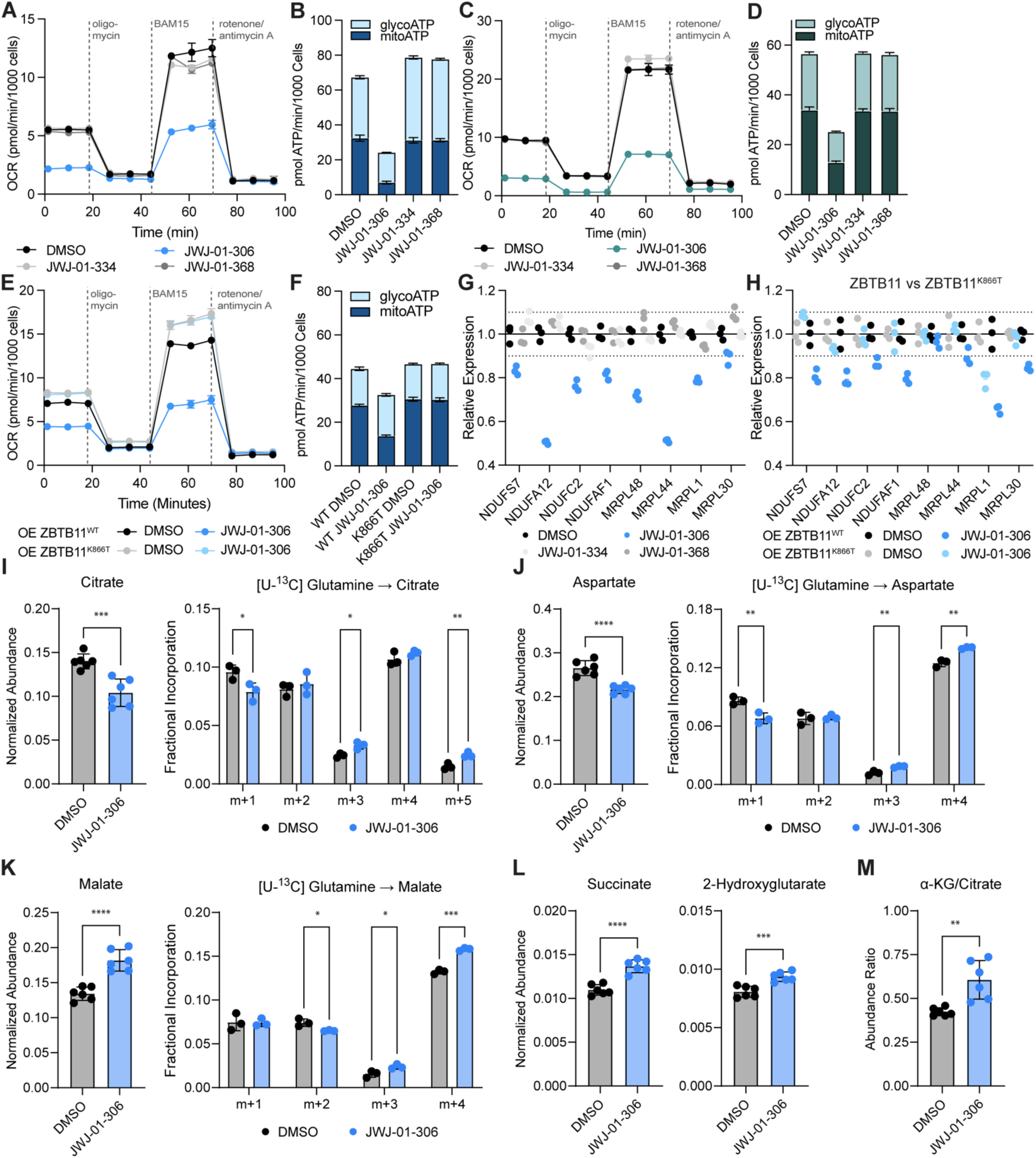
| Metabolic reprogramming of K-Ras inhibitor resistant PDAC cells by ZBTB11 degradation. A. ZBTB11 degradation reduces rates of OXPHOS in sotorasib-resistant MIA PaCa-2 cells. Cells were treated with 10 µM of the indicated compound for 24 hrs. B. ZBTB11 degradation reduces ATP production by OXPHOS in sotorasib-resistant MIA PaCa-2 cells. C. ZBTB11 degradation reduces rates of OXPHOS in MRTX1133-resistant SUIT2 cells. Cells were treated with 10 µM of the indicated compound for 24 hrs. D. ZBTB11 degradation reduces ATP production by OXPHOS in MRTX1133-resistant SUIT2 cells. E. Overexpression of ZBTB11^WT^ or ZBTB11 ^K866T^ rescues JWJ-01-306-mediated reduction of OXPHOS. Sotorasib-resistant MIA PaCa-2 cell lines with stable expression of NanoLuc-ZBTB11^WT^ or NanoLuc-ZBTB11^K866T^ were generated. Cells were treated with 10 µM JWJ-01-306 for 24 hrs. F. Overexpression of ZBTB11^WT^ or ZBTB11 ^K866T^ rescues JWJ-01-306-mediated reduction of ATP production by OXPHOS. G. ZBTB11 degradation induces downregulation of ZBTB11-regulated genes. Sotorasib-resistant MIA PaCa-2 cells were treated with 10 µM of the indicated compound for 24 hrs. H. Overexpression of ZBTB11^WT^ or ZBTB11 ^K866T^ rescues JWJ-01-306-mediated downregulation of ZBTB11-regulated genes. Cells were treated with 10 µM JWJ-01-306 for 24 hrs. I-M. JWJ-01-306 disrupts TCA cycle flux and induces reductive glutamine metabolism. Sotorasib-resistant MIA PaCa-2 cells were treated with DMSO or 10 µM JWJ-01-306 for 24 hrs. Cells were then labeled with ^13^C-labeled glucose or glutamine for an additional 24 hrs followed by global metabolomics analysis, *n* = 3 biological replicates per condition. A-F. Cellular respiration rates were evaluated in a mitochondrial stress test with the indicated drugs using a Seahorse analyzer. OCR and ECAR from the mitochondrial stress test were used to calculate ATP production rates. A-D. Data is depicted as the average +/− S.D. of *n* = 2 biological replicates with *n* = 3 technical replicates each. E-F. Data is depicted as the average +/− S.D. of *n* = 5 biological replicates with *n* = 3 technical replicates each. G-H. mRNA levels were quantified by RT-qPCR. Data is depicted as the average +/− S.D. of *n* = 3 biological replicates and is normalized to DMSO vehicle treatment controls. I-M. Metabolite abundance data is depicted as the average +/− S.D. of *n* = 6 biological replicates (3 replicates each from ^13^C-labeled glucose and glutamine). Mass isotopomer distribution data is depicted as the average +/− S.D. of *n* = 3 biological replicates. Significance level is marked with asterisks (two-tailed student’s t-test, * p-value < 0.05, ** p-value < 0.01, *** p-value < 0.001).

To identify how the expression of mitoribosome and complex I genes downstream of ZBTB11 are directly altered by ZBTB11 degraders to affect the observed proteomic and metabolic changes, we performed qPCR in parental and sotorasib-resistant MIA PaCa-2 cells treated with 10 µM JWJ-01-306, JWJ-01-334 or JWJ-01-368 for 24 hrs. Here, we observed downregulation of transcripts by JWJ-01-306 but not negative controls JWJ-01-334 or JWJ-01-368 (Fig. 4G, Fig. S8C). We confirmed these results in matched experiments in parental and MRTX1133-resistant SUIT2 cells (Fig. S8F-G). To confirm these effects are specific to ZBTB11 degradation, we treated sotorasib-resistant MIA PaCa-2 cells that stably expressed ZBTB11^WT^ or ZBTB11^K866T^ with 10 µM JWJ-01-306 for 24 hrs and assessed levels of relevant downstream transcripts by qPCR. Here, we observed complete rescue of transcript loss by the degradation-resistant ZBTB11^K866T^ mutant for 7/8 downstream genes, and partial rescue of MRPL1 (Fig. 4H). Finally, we examined effects of ZBTB11 degraders on ZBTB11 and K-Ras transcript levels, where we observed compensatory upregulation of ZBTB11 in the presence of JWJ-01-306 but not negative controls JWJ-01-334 and JWJ-01-368 in parental and resistant MIA PaCa-2 cells and SUIT2 cells (Fig. S8C, F, G, H). Varied effects on K-Ras transcript levels were observed, indicating that K-Ras transcription is not regulated by ZBTB11 (Fig. S8C, F, G, H).

To identify the metabolic pathway alterations that mediate the effects of ZBTB11 degradation on cellular respiration, we performed metabolomics on sotorasib-resistant MIA PaCa-2 cells treated with 10 µM JWJ-01-306 for 24 hrs and labeled with ^13^C-glucose or ^13^C-glutamine. Metabolite set enrichment analysis revealed JWJ-01-306 treatment induced changes consistent with reduced OXPHOS, including in the TCA cycle and pyruvate metabolism (Fig. S9K, Dataset 5). Manual inspection of metabolite abundances and mass isotopomer distributions aligned with this analysis and indicated significant changes in the TCA cycle pathway. (Fig. 4I-M, Dataset 5). The cellular abundance of citrate, a TCA cycle intermediate, was reduced with JWJ-01-306 treatment (Fig. 4I). As tumor cells utilize reductive glutamine metabolism to support various cellular processes when activity of the electron transport chain (ETC) is inhibited or the TCA cycle is perturbed^47,48^, we sought to determine whether ZBTB11 degradation would promote reductive glutamine metabolism. We found that the labeling pattern of citrate (derived from ^13^C-glutamine) in cells treated with JWJ-01-306 was altered, with an increased proportion of m+3 and m+5 isotopomers, indicative of increased reductive carboxylation of glutamine-derived α-ketoglutarate (Fig. 4I). Consistent with these results, we also observed JWJ-01-306-dependent increases in aspartate, malate and fumarate m+3 isotopomers indicative of reductive glutamine metabolism (Fig. 4J-K, Dataset 5). Furthermore, shifts in TCA substrate abundances (Fig. 4I-4L) and the α-ketoglutarate/citrate ratio (Fig. 4M) were in agreement with previous observations in cells with perturbed mitochondrial respiration^49,50^. Taken together, these data demonstrate that JWJ-01-306 treatment impairs the ETC and disrupts normal flux through the TCA cycle.

### ZBTB11 degraders show reduced neurotoxicity compared to complex I inhibitors

Having established the mechanism-of-action of JWJ-01-306, we next sought to compare ZBTB11 degradation to complex I inhibition in human models of neuronal health. The primary safety liability of OXPHOS-targeting drugs are toxic effects on neurons, which rely heavily on mitochondrial metabolism to sustain their bioenergetic needs.^27^ These effects often go unnoticed in murine studies, due to species differences between the human and murine nervous system, and challenges with accurately measuring neuropathy in mice.^51^ Furthermore, molecular glue degraders often have different target profiles in murine and human systems due to sequence differences in both CRBN and the targets that alter the induced protein-protein interaction interface^52–54^. To overcome these limitations and provide an early readout of the propensity of our ZBTB11 degraders to cause neuropathy, we profiled the neurotoxicity of ZBTB11 degraders using a suite of assays to monitor neuronal viability, morphology, and mitochondrial function in human induced pluripotent stem cell (hiPSC)-derived neurons.^55–58^ We found that hiPSC-derived neurons exposed to 1 µM IACS-010759 for 24 hrs exhibit significantly decreased mitochondrial membrane potential (MMP) and increased reactive oxygen species (ROS), consistent with inhibition of mitochondrial complex I and reported adverse events in clinical trials^27^ (Fig. 5A). Following treatment with 1 µM IACS-010759 for 72 hrs, neurite length and neuronal survival were also decreased (Fig. 5B). In contrast, 1 µM JWJ-01-306 showed minimal effects on all phenotypes assayed (Fig. 5A-B). Together, these results show our panel accurately identifies neurotoxic compounds targeting mitochondrial function and demonstrate that ZBTB11 degradation has reduced acute neuronal toxicity *in vitro* relative to complex I inhibition.

**Figure 5.**
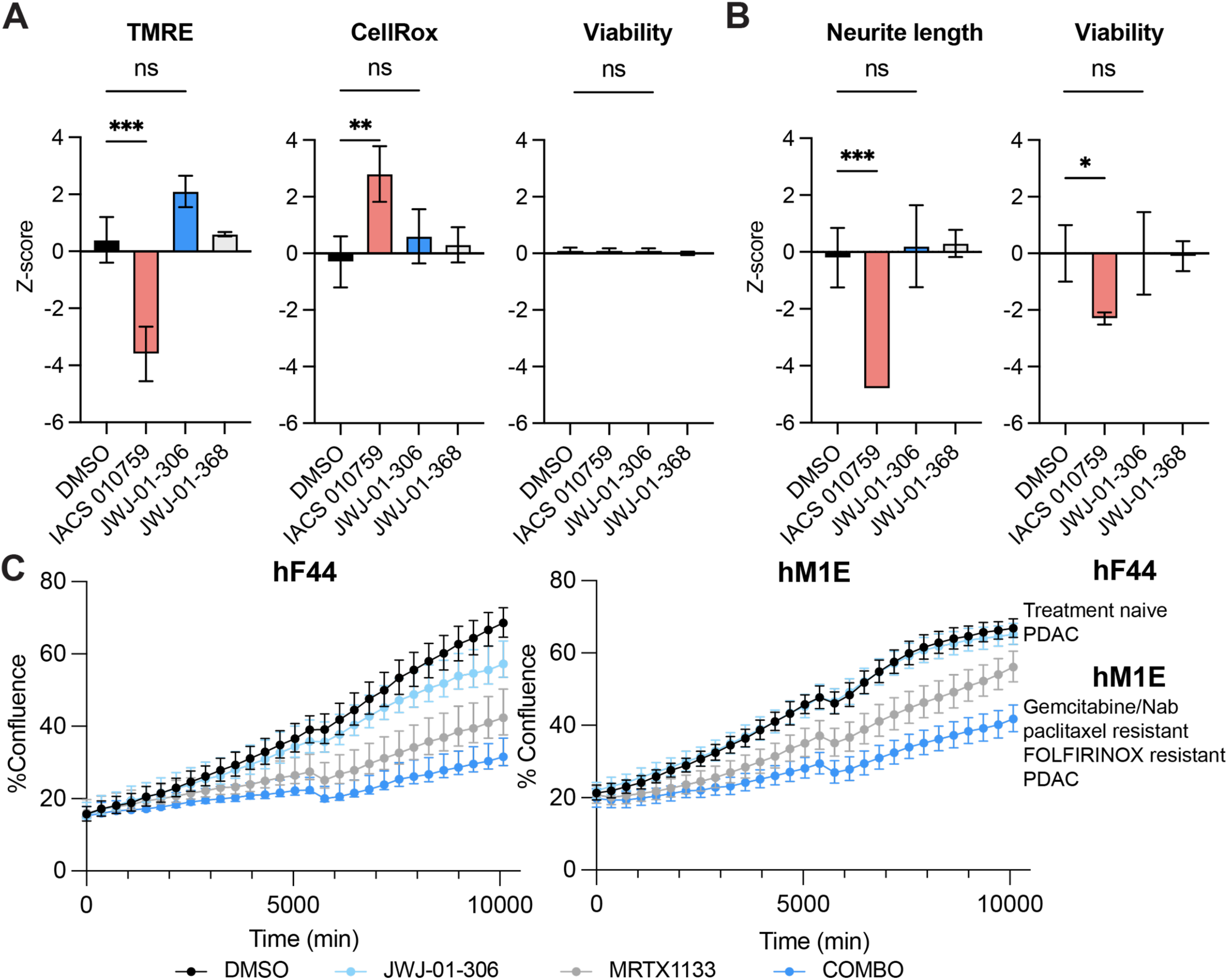
| ZBTB11 degradation spares neurons and deepens the response of PDAC patient-derived organoids to K-Ras inhibitors. A. Mitochondrial activity and oxidative stress assays show differential pharmacology of Complex I and ZBTB11 perturbation. hiPSC-derived neurons were treated for 24 h with DMSO, 1 µM IACS 010759, 1 µM JWJ-01-306 or 1 µM JWJ-01-368. B. Neurotoxicity assays show differential pharmacology of Complex I and ZBTB11 perturbation. hiPSC-derived neurons were treated for 72 h with DMSO, 1 µM IACS 010759, 1 µM JWJ-01-306 or 1 µM JWJ-01-368. C. Proliferation assays show ZBTB11 + K-Ras inhibitor combination has superior antiproliferative effects in PDAC patient-derived organoids. Cells were treated with DMSO vehicle, 1 µM JWJ-01-306, 200 nM MRTX1133 or 1 µM JWJ-01-306 + 200 nM MRTX1133 (COMBO) and cellular confluence observed using brightfield imaging at the indicated timepoints. A.C. Data plotted as mean +/− S.D. of *n =* 3 biological replicates.

### ZBTB11 degraders deepen the response to K-Ras inhibition in PDAC patient-derived organoids

To validate ZBTB11 degrader co-treatment as a potential therapeutic strategy in a more clinically relevant context, we turned to patient-derived organoid models of K-Ras^G12D^ driven PDAC^59^. Personalized tumor drug sensitivity profiling using these models was shown to correlate with patient-responses, and their predictive value is currently being assessed in a clinical trial (NCT04469556). As MRTX1133 is still being evaluated in early phase clinical trials, no MRTX1133-resistant patient-derived organoids are available. We therefore sought to evaluate if JWJ-01-306 could deepen the response to MRTX1133 as a combination therapy in organoids derived from tumor samples from both a chemotherapy-naive patient (hF44) and a highly pretreated multi-drug resistant patient (hM1E, Fig. 5C). We treated organoids with DMSO, JWJ-01-306 (1 µM), MRTX1133 (100 nM), or JWJ-01-306 (1 µM) + MRTX1133 (100 nM) combination and monitored cell proliferation by brightfield imaging over 7 d (Fig. 5C). Both organoids had a partial response to MRTX1133, and greater anti-proliferative effects were achieved by combining treatment with JWJ-01-306, indicating that enhanced responses to K-Ras inhibitors can be accomplished by preemptive blockade of metabolic escape pathways. Together, these experiments establish ZBTB11 degradation as an attractive therapeutic strategy to combat the aberrant metabolic switch to OXPHOS that drives resistance to K-Ras inhibitors.

## DISCUSSION

In this manuscript, we disclose molecular glue degraders of the transcription factor ZBTB11. By generating a comprehensive set of chemical and genetic tools we identified ZBTB11-governed metabolic networks in human PDAC. In contrast to existing OXPHOS inhibitors, most of which are thought to target complex I-V enzymatic activity, JWJ-01-306 directly targets the transcriptional upregulation of OXPHOS genes associated with cancer drug resistance. We demonstrate that pharmacological degradation of ZBTB11 counters this metabolic reprogramming associated with acquired resistance to K-Ras inhibitors in multiple *ex vivo* models of PDAC, identifying ZBTB11 as a druggable therapeutic vulnerability in PDAC.

A major limitation for targeting aberrant OXPHOS are the neurotoxic side effects of complex I inhibition, observed as on-target adverse events in clinical trials and in follow-up murine studies.^27^ Using human iPSC-derived neuronal models, we demonstrated that ZBTB11 degradation had minimal effects on neuronal viability and function at phenotypically relevant time points (3d), in contrast to complex I inhibitor IACS-010759. While comprehensive studies in *in vivo* mammalian models at extended time points will be critical for establishing the preclinical therapeutic window of ZBTB11 degradation in PDAC, our work marks a critical milestone by identifying a mechanistically orthogonal therapeutic strategy for targeting high OXPHOS cancer states.

Our patient-derived organoid study demonstrated that concurrent K-Ras and ZBTB11 blockade deepens responses to K-Ras^G12D^ inhibition in PDAC organoids that had not been previously treated with MRTX1133, including in the multi-drug resistant PDAC line hM1E. ZBTB11 degradation may therefore have applications in other cancers that upregulate OXPHOS to drive resistance to therapeutic interventions, including chemotherapies^60–62^, anti-angiogenic therapies^63^, and targeted therapies such as inhibition of BCL2^64^, tyrosine kinases^65^, and BRAF^66^. We anticipate that our suite of chemical and genetic tools will accelerate efforts to map ZBTB11 sensitive cancer cell states and identify additional tumor types where ZBTB11 depletion can address cancer drug resistance.

## ACKNOWLEDGEMENTS

N.L.T. was supported by the NSF Graduate Research Fellowship (DGE-2038238) and the NIH/NCI Cancer Cell Signaling & Communication Training Grant (T32CA009523). G.E.G. was supported by an NIH Molecular Biophysics Training Grant (T32GM139795). Research reported in this publication was supported by the NCI Cancer Center Support Grant P30 CA030199. K.C.M.G., H.T. and A.M.L. were supported by a grant from The Lustgarten Foundation and the Research for a Cure of Pancreatic Cancer fund. We would like to acknowledge support from the UCSD BMPMSF Proteomics Core, and the SBP Proteomics, Cancer Metabolism, and Functional Genomics Core Facilities. We would like to acknowledge Nathanael Gray for providing the CRBN-targeting molecular glue screening library.

## AUTHOR CONTRIBUTIONS

N.L.T. performed Seahorse, RT-qPCR, HiBiT and NanoLuc luminescence assays, NanoBRET ternary complex assays, trypan blue exclusion, computational modeling, generated inducible CRISPRi cell lines, and prepared samples for proteomics and metabolomics studies. N.L.T. and M.M. conducted the proteomics experiments, performed proteomics and metabolomics analysis, and prepared figures. J.W.J. designed and performed molecule synthesis. C.H. designed and generated CRISPRi constructs. D.A.S. performed metabolomic studies in collaboration with N.L.T. G.E.G performed CRBN target engagement assay. N.L.T and E.S.W. designed and cloned DNA plasmid constructs, generated ZBTB11-HiBiT cell lines, and performed immunoblot experiments. E.S.W supervised the research, performed study design and analysis, prepared figures, and edited the manuscript. F.M.F. supervised the research, performed study design and analysis, funding acquisition, prepared the figures, wrote the manuscript (first draft). K.C.M.G., H.T. and A.M.L. developed MRTX1133-resistant cell lines, performed bright-field proliferation assays and edited the manuscript. All authors have read and approved the final manuscript.

---

**Supporting Figure 1.**
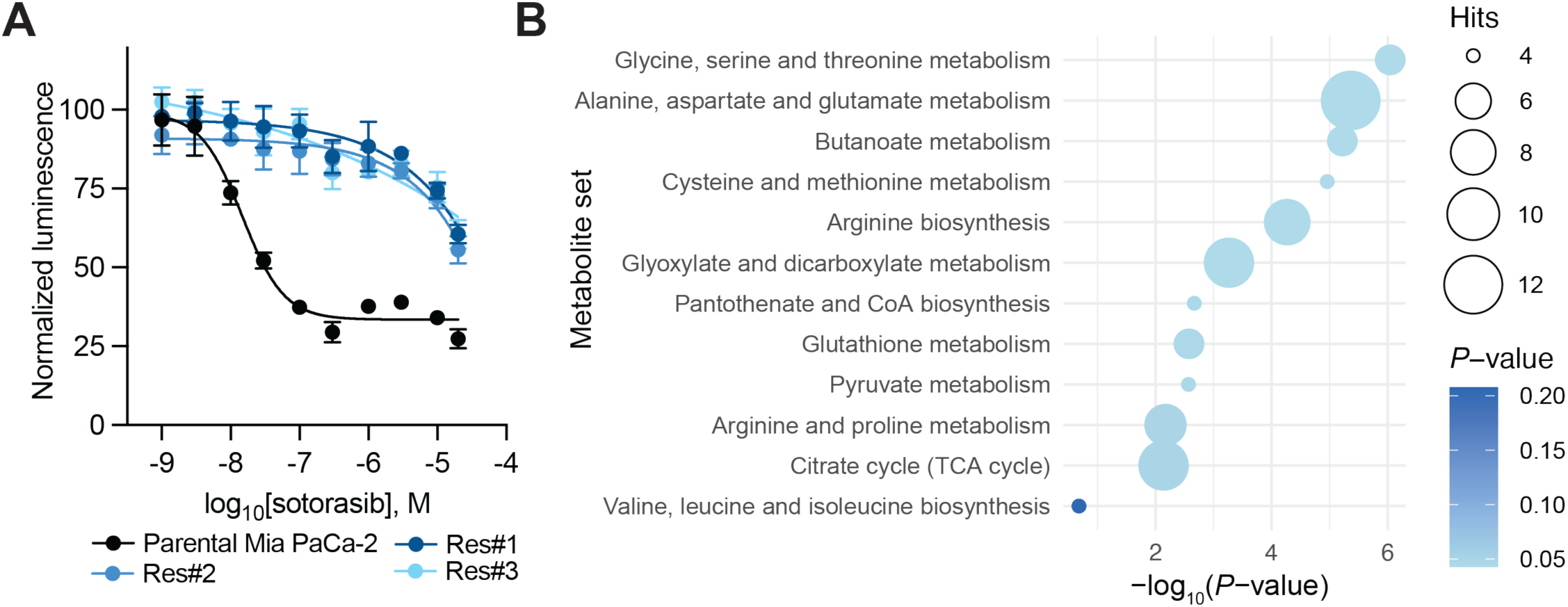
| Metabolomic profiling of parental and K-Ras G12C inhibitor resistant PDAC cell lines. A. Sotorasib/AMG-510-resistant MIA PaCa-2 cells are 100-fold less sensitive to sotorasib/AMG-510 than parental MIA PaCa-2 cells. Cells were treated with the indicated dose of sotorasib for 72 hrs, and viability was evaluated by CellTiterGlo®. Data is depicted as the average +/− standard deviation (S.D.) of *n* = 3 biological replicates and is normalized to DMSO vehicle treatment controls. B. MSEA identifies amino acid metabolic pathways as major metabolic shifts between parental and sotorasib-resistant MIA PaCa-2 cells. Parental and sotorasib-resistant MIA PaCa-2 cells were labeled with ^13^C-labeled glucose or glutamine for 24 hrs followed by global metabolomics analysis, *n* = 3 biological replicates per condition. Metabolite abundance changes were analyzed by the global test (*P* < 0.05). Metabolites were then evaluated using metabolite set enrichment analysis (MSEA) using KEGG pathways and filtered for pathways containing a minimum of four hits. Full datasets in Dataset 1.

**Supporting Figure 2.**
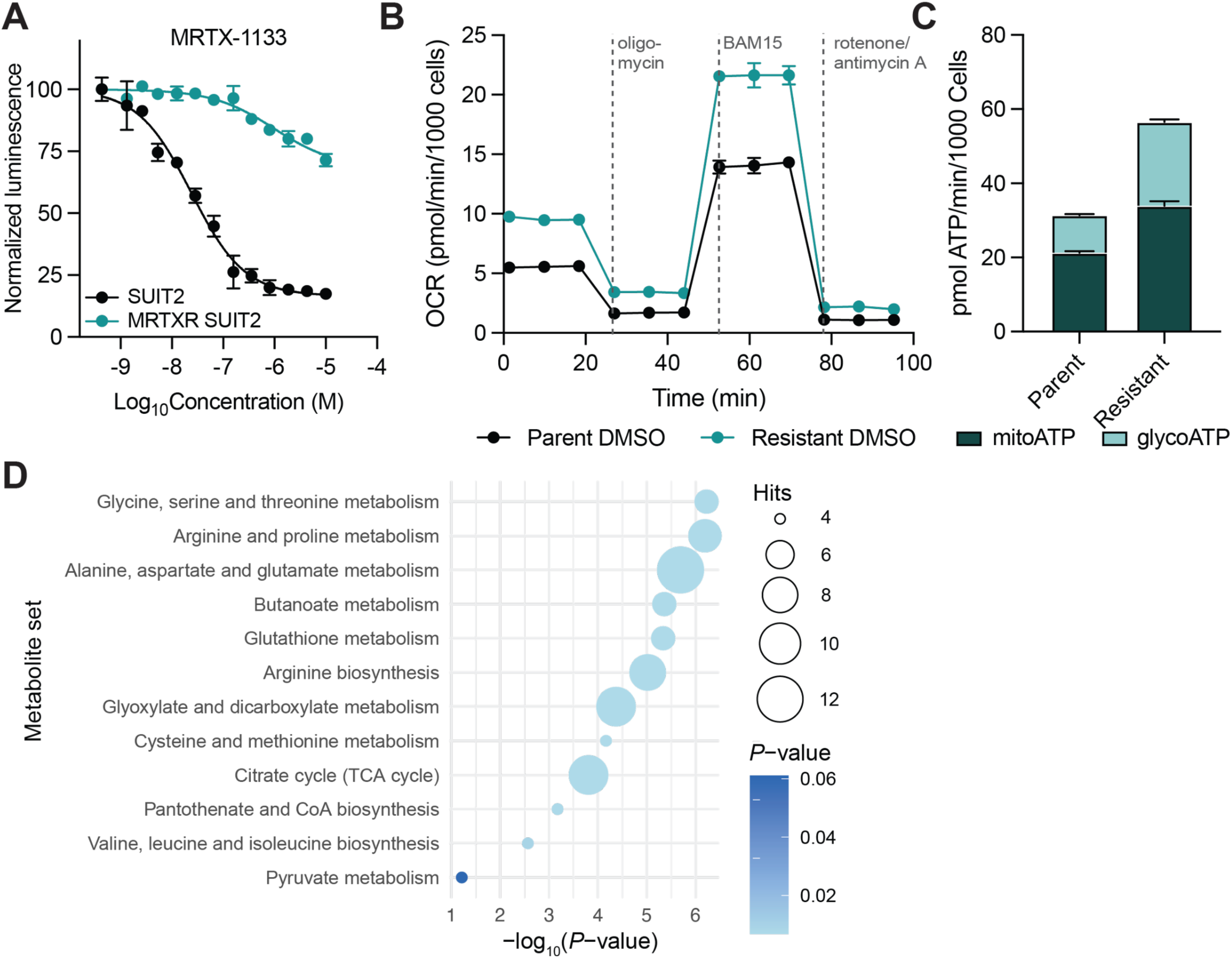
| Metabolic profiling of parental and K-Ras G12D inhibitor resistant PDAC cell lines. A. MRTX1133-resistant SUIT2 cells are 100-fold less sensitive to MRTX1133 than parental SUIT2 cells. Cells were treated with the indicated dose of MRTX1133 for 72 hrs, and viability was evaluated by CellTiterGlo®. Data is depicted as the average +/− standard deviation (S.D.) of *n* = 3 biological replicates and is normalized to DMSO vehicle treatment controls. B. MRTX1133-resistant SUIT2 cells perform higher basal and maximal levels of OXPHOS than parental SUIT2 cells. C. MRTX1133-resistant SUIT2 cells are more energetic than parental SUIT2 cells. B-C. Cellular respiration rates were evaluated in a mitochondrial stress test with the indicated drugs using a Seahorse analyzer. OCR and ECAR from the mitochondrial stress test were used to calculate ATP production rates. Data is depicted as the average +/− S.D. of *n* = 2 biological replicates with *n* = 3 technical replicates each. D. MSEA identifies oxidative stress and amino acid-related pathways as major metabolic shifts between parental and MRTX1133-resistant SUIT2 cells. Parental and MRTX1133-resistant SUIT2 cells were labeled with ^13^C-labeled glucose or glutamine for 24 hrs followed by global metabolomics analysis, *n* = 3 biological replicates per condition. Metabolite abundance changes were analyzed by the global test (*P* < 0.05). Metabolites were then evaluated for pathway enrichment using KEGG pathway analysis and filtered for pathways containing a minimum of four hits. Full datasets in Dataset 1.

**Supporting Figure 3.**
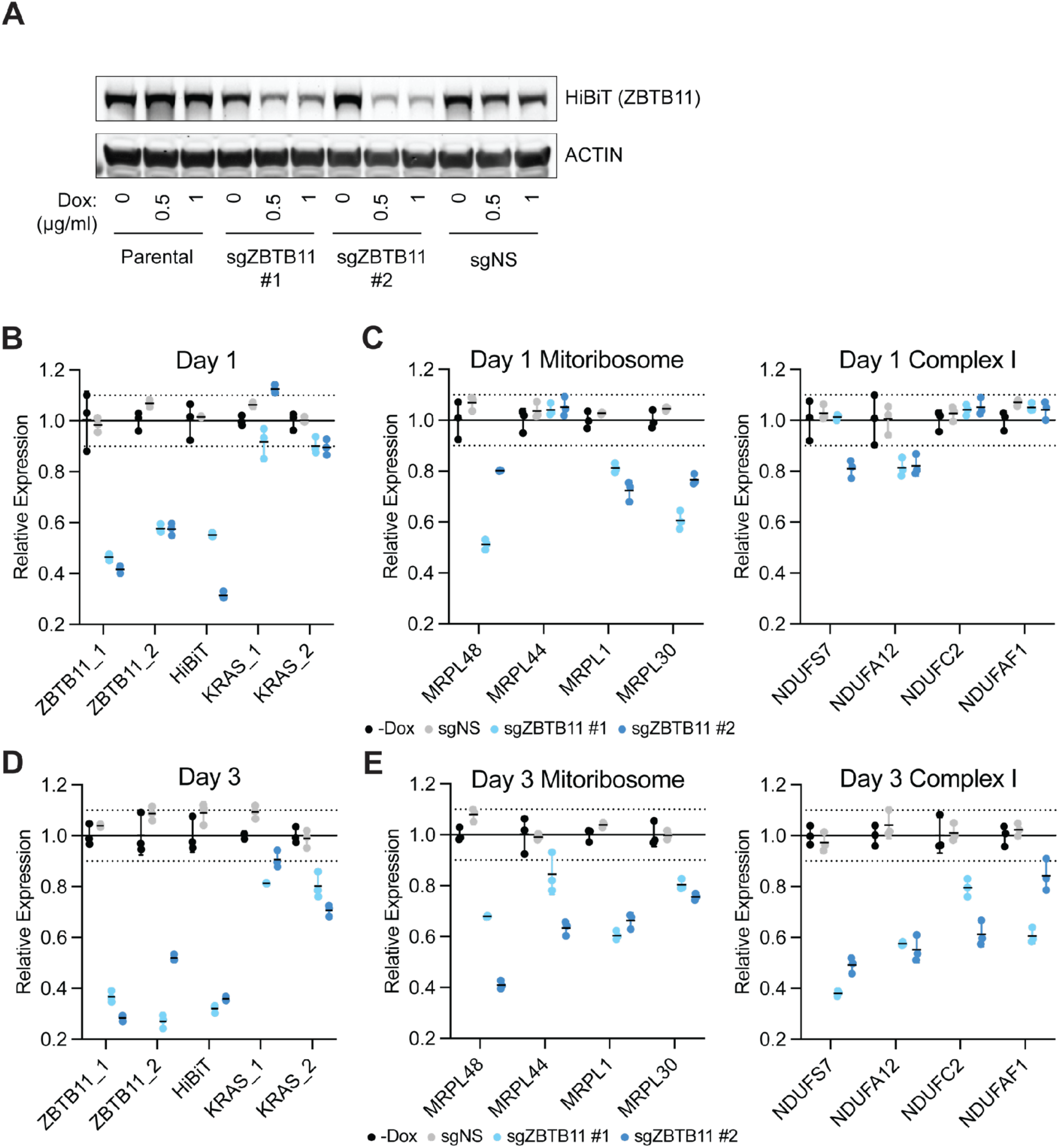
| CRISPRi ZBTB11 knockdown impacts OXPHOS-related genes. A. ZBTB11-targeting sgRNAs facilitate CRISPRi-mediated knockdown of ZBTB11. Cells were treated with 500 ng/ml of doxycycline for 72 hrs to induce Cas9 expression, and protein levels were quantified by WB. Depicted blots are representative of *n* = 2 independent experiments. Uncropped blots found in Source Data. B-E. CRISPRi-mediated knockdown induces downregulation of ZBTB11 and ZBTB11-regulated genes. Cells were treated with 500 ng/ml of doxycycline for 24 or 72 hrs to induce Cas9 expression, and mRNA levels were quantified by RT-qPCR. Data is depicted as the average +/− S.D. of *n* = 3 biological replicates and is normalized to H_2_O vehicle treatment controls.

**Supporting Figure 4.**
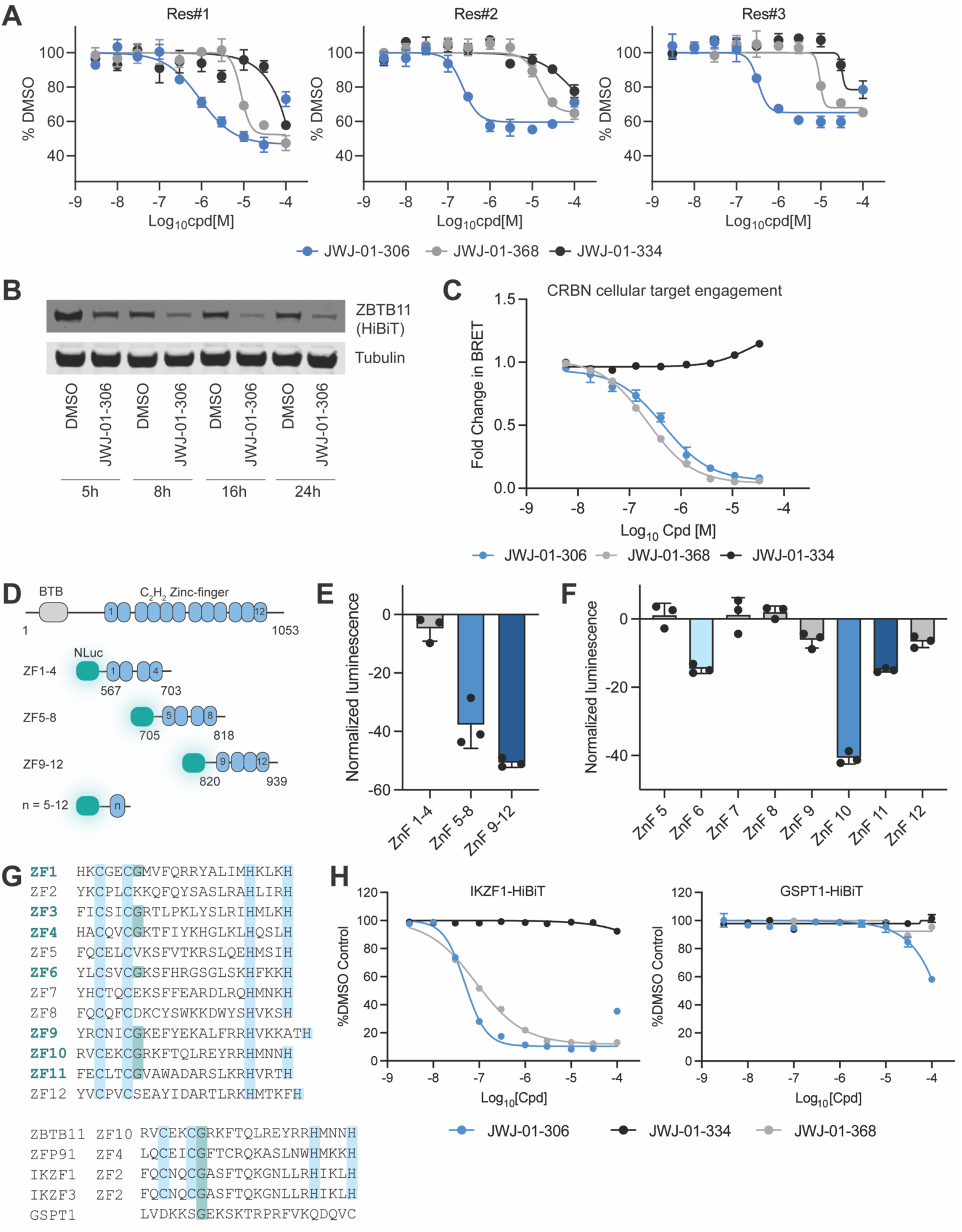
| Characterization of the ZBTB11 degrader pharmacology. A. JWJ-01-306, but not negative controls, degrades ZBTB11-HiBiT in sotorasib-resistant MIA PaCa-2 knock-in cells. Cells were treated for 5 hrs with the indicated compound. B. Time-course ZBTB11-HiBiT degradation. Cells were treated with 10 µM JWJ-01-306 for indicated amount of time, and protein levels were quantified by western blot. Depicted blots are representative of *n* = 2 independent experiments. Uncropped blots found in Source Data. C. Cellular CRBN engagement. HEK293 cells were transiently transfected with NanoLuc-CRBN and treated with the NanoBRET™ Tracer Reagent and indicated compound for 2 hrs. D. Construct design for ZBTB11 ZF fingerprinting experiments. E-F. ZBTB11 ZF fingerprinting reveals ZF10 as the primary CRBN target degron. MOLT-4 cell lines with stable expression of the indicated constructs were generated. Cells were treated with 1 µM screening hit compound ALV-05-184 for 5 hrs. G. Sequence alignment of the ZBTB11 ZF domains and common IMiD target proteins with C_2_H_2_, and CXXCG residues highlighted. H. IKZF1-HiBiT and GSPT1-HiBiT degradation in Jurkat knock-in cells. Cells were treated for 8 hrs with the indicated compound. A, C, E-F, H. Data is depicted as the average +/− S.D. of *n* = 3 biological replicates and is normalized to DMSO vehicle treatment control.

**Supporting Figure 5.**
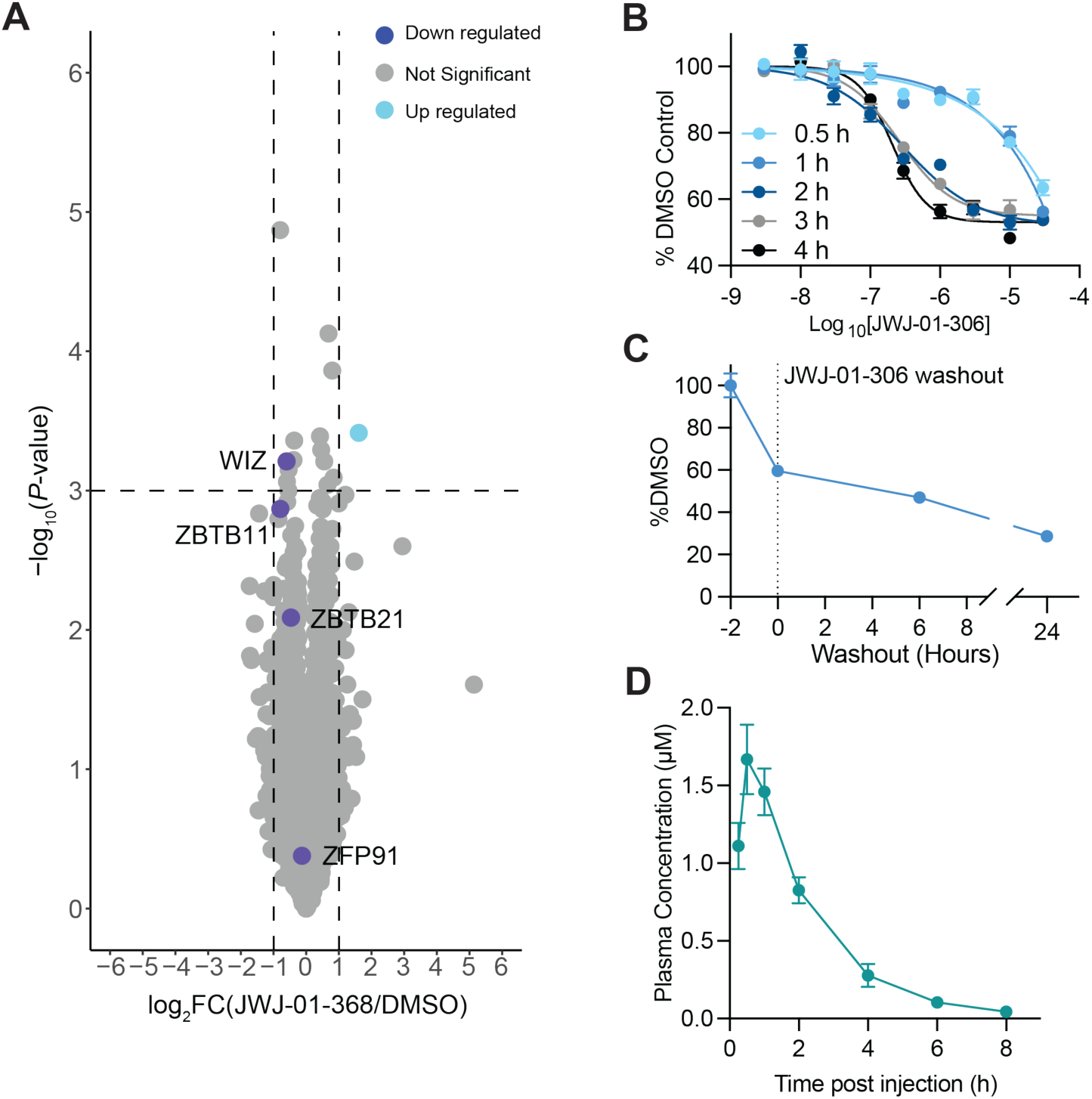
| Further characterization of ZBTB11 degrader compound and controls. A. Global proteomics analysis of MIA PaCa-2 cells treated with 10 µM JWJ-01-368 for 5 hrs. Samples were prepared as *n* = 3 biological replicates. Full datasets in Dataset 2. B. ZBTB11-HiBiT degradation by JWJ-01-306 achieves optimal kinetics by 2 hrs. MIA PaCa-2 knock-in cells were treated for the indicated time. C. ZBTB11-HiBiT protein levels decrease following JWJ-01-306 washout. MIA PaCa-2 knock-in cells were treated for 2 hrs with 10 µM JWJ-01-306, followed by three media washes to remove drug. ZBTB11 levels were evaluated by ZBTB11-HiBiT luminescence assay. D. Pharmacokinetics of JWJ-01-306 in C57BL/6 mice dosed intraperitoneal with 10 mg/kg JWJ-01-306-HCl. Full datasets in Dataset 3. B-C. Data is depicted as the average +/− S.D. of *n* = 3 biological replicates and is normalized to DMSO vehicle treatment control.

**Supporting Figure 6.**
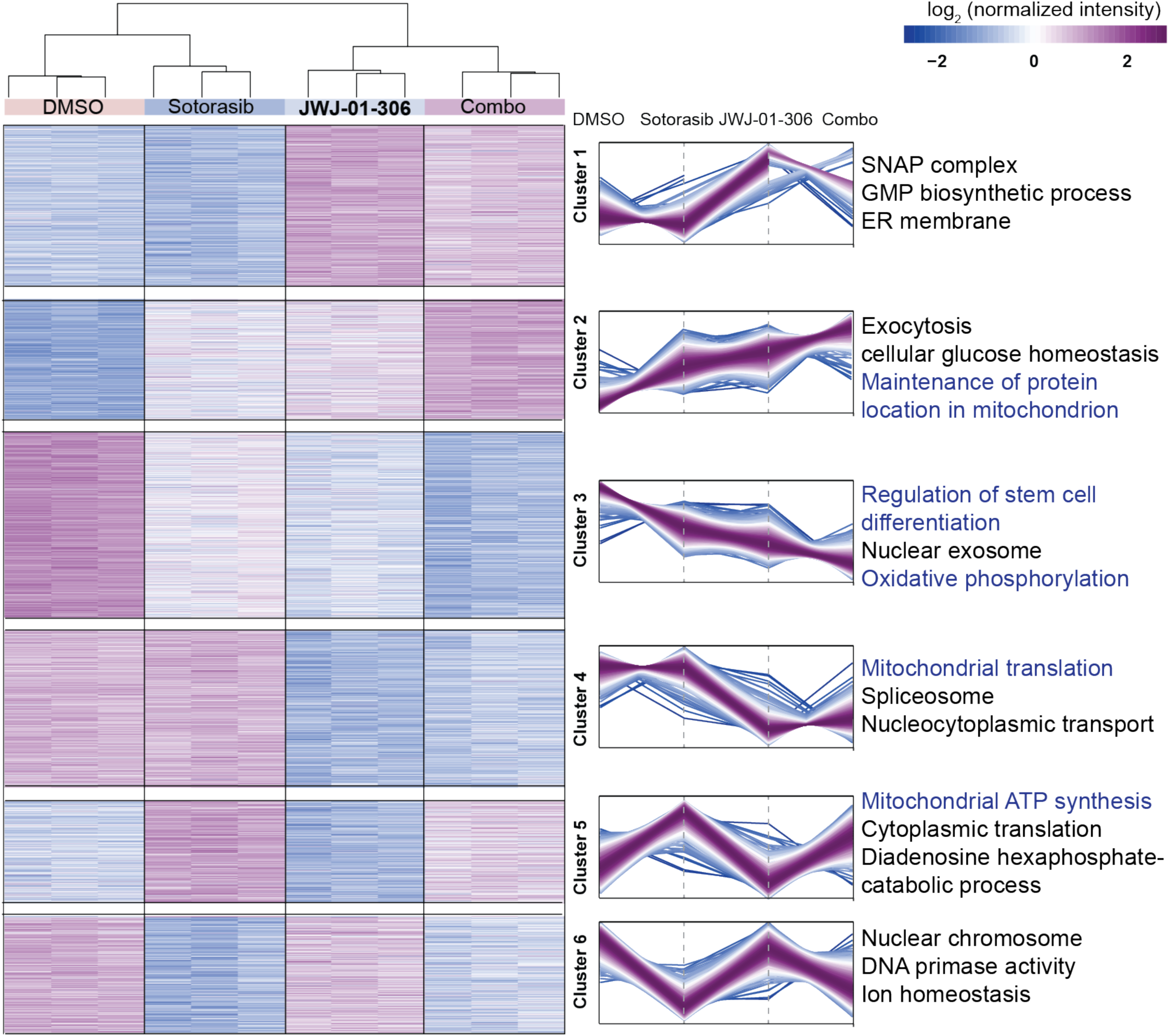
| Cellular processes targeted by K-Rasi+ZBTB11 degrader combinations in K-Rasi resistant PDAC at 5d. Sotorasib-resistant MIA PaCa-2 cells were treated with DMSO, 1 µM sotorasib, 10 µM JWJ-01-306, or 1 µM sotorasib + 10 µM JWJ-01-306 (combo) for 120 hrs followed by global proteomics analysis, *n =* 3 biological replicates per condition. Protein abundance changes were analyzed by one-way ANOVA test (FDR < 0.05) and clustered using K-nearest neighbors clustering. Each line within the clusters is color-coded according to its distance from the cluster center, ranging from purple (close) to light blue (far). Proteins in each cluster were then evaluated for pathway enrichment using GO and KEGG pathway analysis. Representative pathways are shown and pathways associated with ZBTB11 function are highlighted in blue text. Full datasets in Dataset 4.

**Supporting Figure 7.**
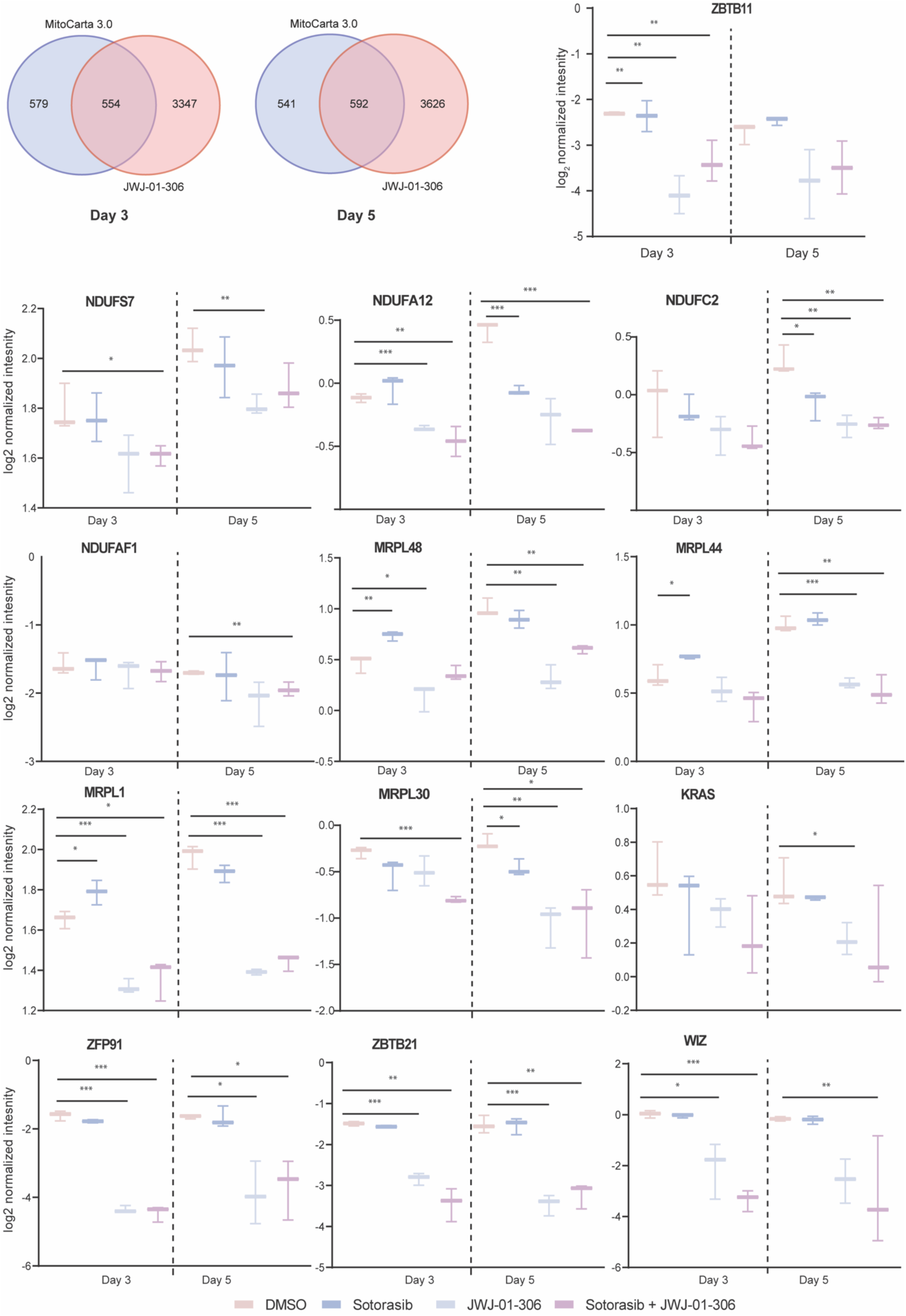
| JWJ-01-306 leads to reduction in ZBTB11 downstream mitochondrial proteins. A. Comparison between MitoCarta 3.0 database and significantly regulated protein hits identified in proteomics reveals enrichment of mitochondria-related proteins. B. Protein abundances of ZBTB11-regulated targets decrease with treatment of 10 µM JWJ-01-306. Each boxplot represents the median value, with the bounds indicating the 25^th^ and 75^th^ percentiles. Significance level is marked with asterisks (two-tailed moderated t-test, * p-value < 0.05, ** p-value < 0.01, *** p-value < 0.001).

**Supporting Figure 8.**
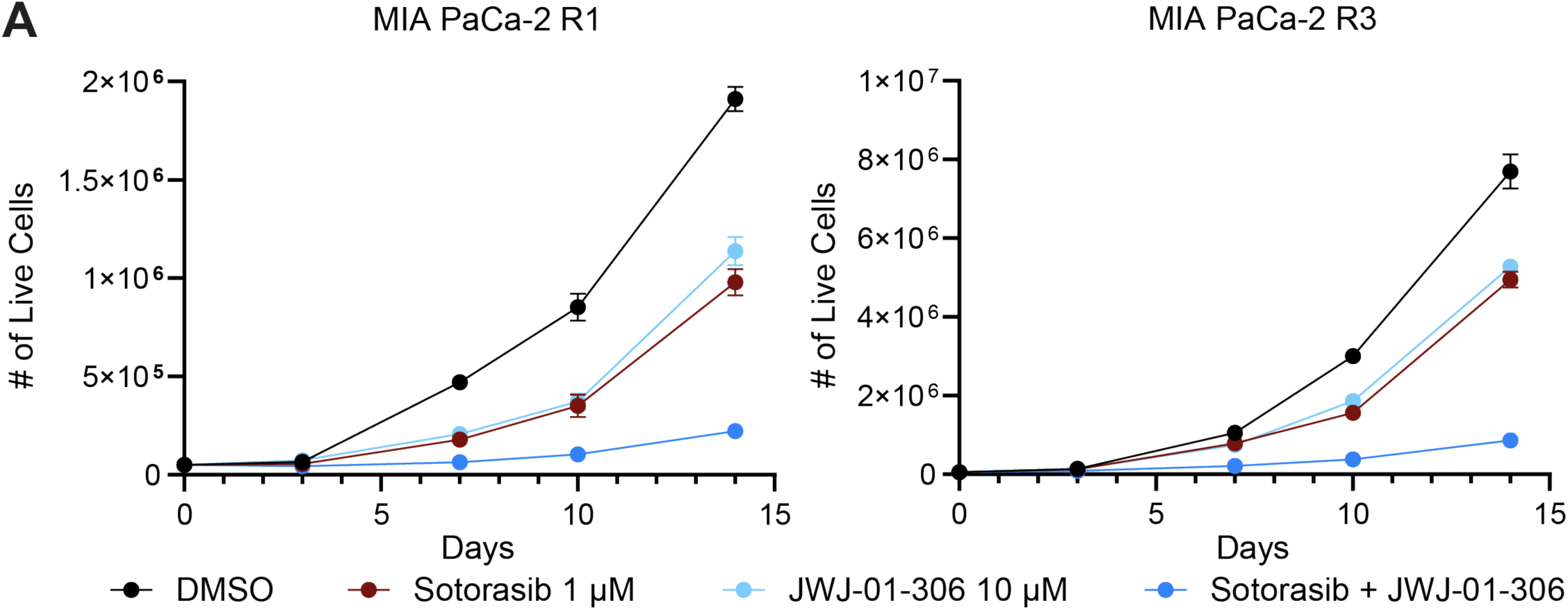
| JWJ-01-306 combines with K-Ras inhibition to overcome acquired resistance. JWJ-01-306 synergizes with sotorasib to inhibit proliferation of sotorasib-resistant MIA PaCa-2 cells. Cells were treated with the indicated compounds and cell numbers were measured using trypan blue exclusion. Data is depicted as the average +/− S.D. of *n* = 3 biological replicates.

**Supporting Figure 9.**
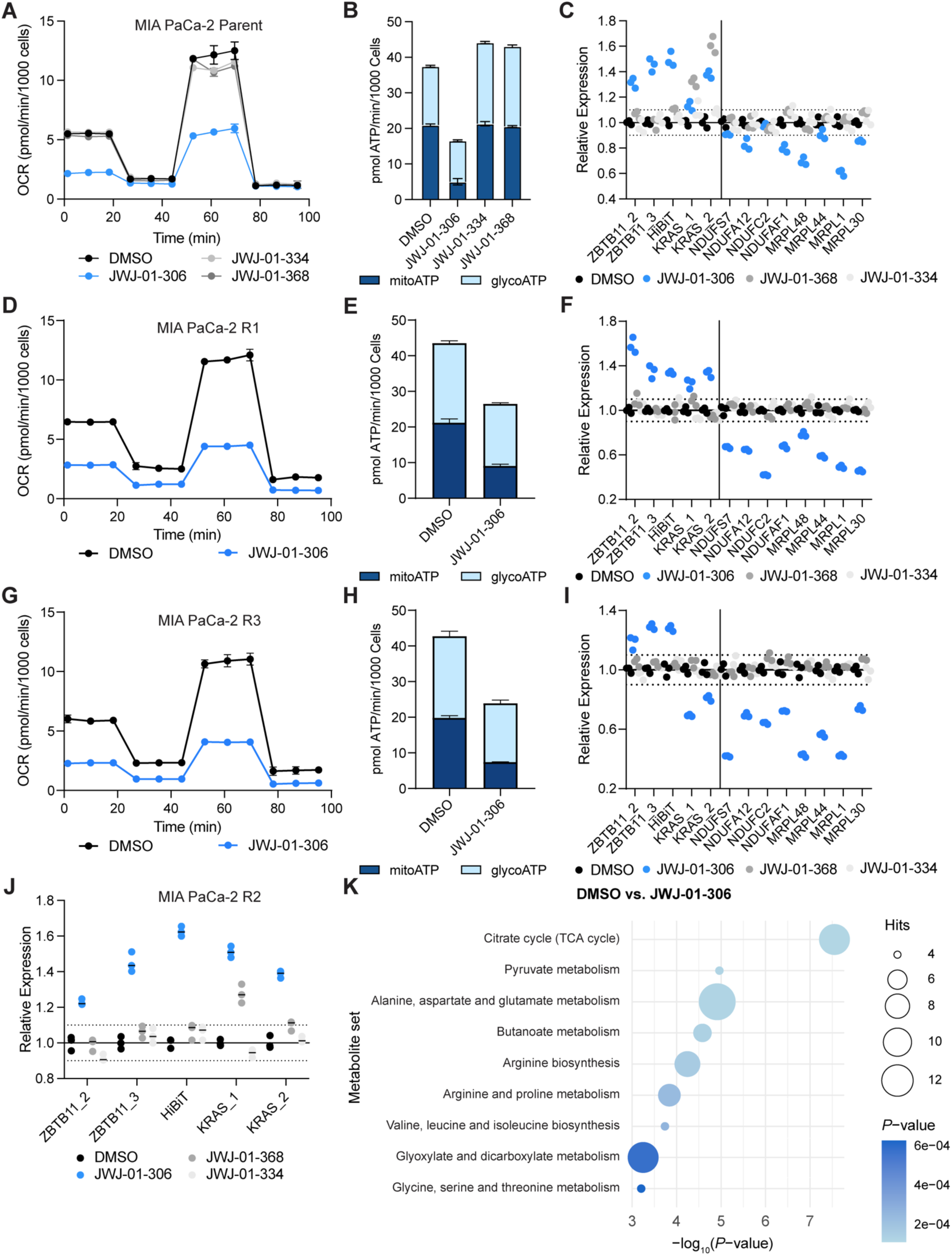
| Metabolic reprogramming of parental and K-Ras^G12C^ inhibitor resistant PDAC cells by ZBTB11 degradation. A-B, D-E, G-H. ZBTB11 degradation reduces rates of OXPHOS in parental and sotorasib-resistant MIA PaCa-2 cells. C, F, I-J. ZBTB11 degradation induces compensatory genetic upregulation of ZBTB11 and downregulation of ZBTB11-regulated genes in MIA PaCa-2 cells. K. MSEA identifies the TCA cycle as the major metabolic pathway disrupted by ZBTB11 degradation. Sotorasib-resistant MIA PaCa-2 cells were treated with DMSO or 10 µM JWJ-01-306 for 24 hrs. Cells were then labeled with ^13^C-labeled glucose or glutamine for an additional 24 hrs followed by global metabolomics analysis, *n* = 3 biological replicates per condition. Metabolite abundance changes were analyzed by the global test (*P* < 0.05). Metabolites were then evaluated for pathway enrichment using KEGG pathway analysis and filtered for pathways containing a minimum of four hits. Full datasets in Dataset 5. A-J. Cells were treated with 10 µM of the indicated compound for 24 hrs. A-B, D-E, G-H. Cellular respiration rates were evaluated in a mitochondrial stress test with the indicated drugs using a Seahorse analyzer. OCR and ECAR from the mitochondrial stress test were used to calculate ATP production rates. C, F, I. mRNA levels were quantified by RT-qPCR. Data is depicted as the average +/− S.D. of *n* = 3 biological replicates and is normalized to DMSO vehicle treatment controls.

**Supporting Figure 10.**
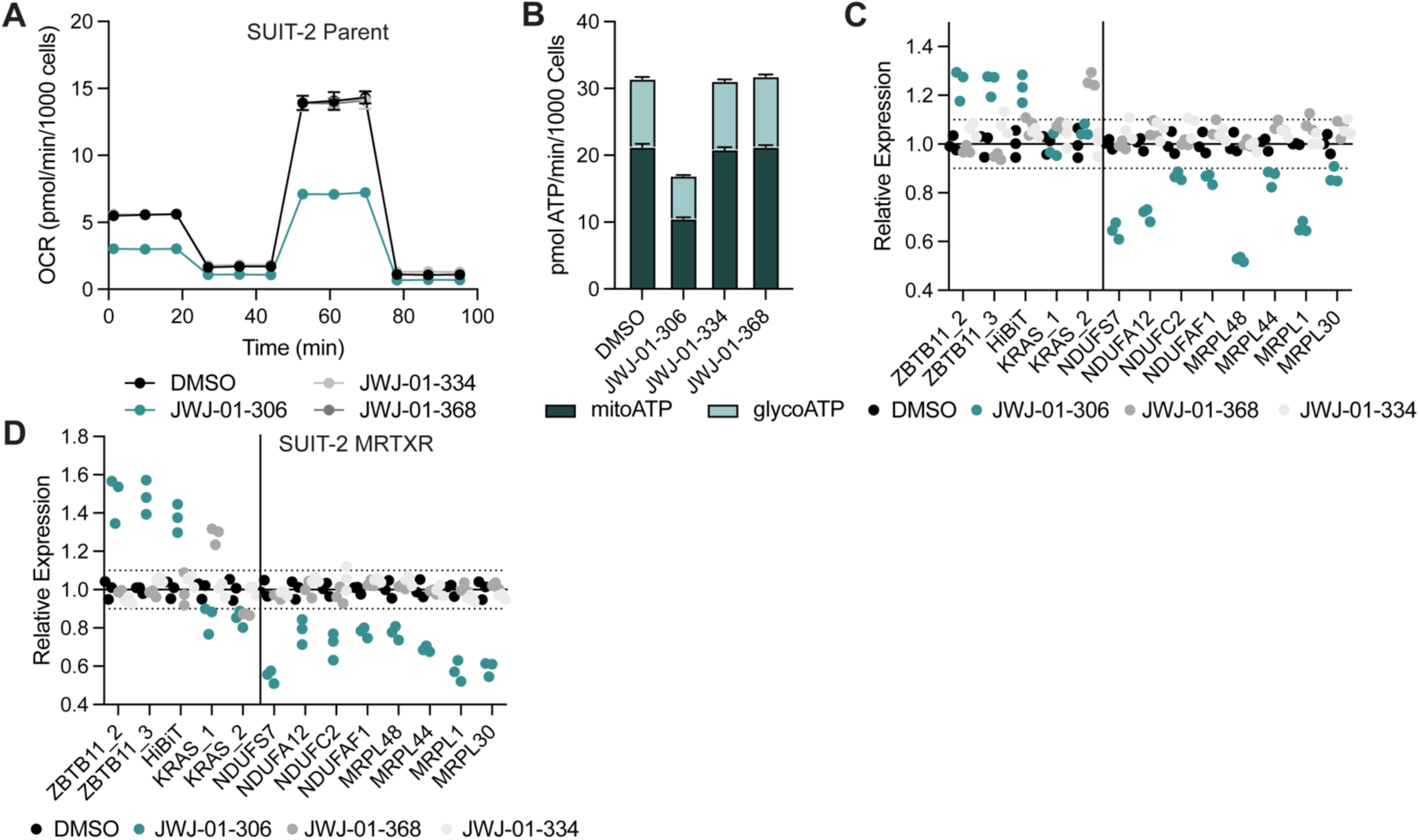
| Metabolic reprogramming of parental and K-Ras^G12D^ inhibitor resistant PDAC cells by ZBTB11 degradation. A-B. ZBTB11 degradation reduces rates of OXPHOS in parental SUIT2 cells. C-D. ZBTB11 degradation induces compensatory genetic upregulation of ZBTB11 and downregulation of ZBTB11-regulated genes in SUIT2 cells. A-D. Cells were treated with 10 µM of the indicated compound for 24 hrs. A-B. Cellular respiration rates were evaluated in a mitochondrial stress test with the indicated drugs using a Seahorse analyzer. OCR and ECAR from the mitochondrial stress test were used to calculate ATP production rates. C-D. mRNA levels were quantified by RT-qPCR. Data is depicted as the average +/− S.D. of *n* = 3 biological replicates and is normalized to DMSO vehicle treatment controls.

